# Image-Guided Injectable Niche for Hepatocyte Transplantation

**DOI:** 10.64898/2026.01.30.702821

**Authors:** Vardhman Kumar, Joa Yun, Susanna K. Elledge, Nicole Henning, Katarzyna A. Grzelak, Ashley D. Westerfield, Amy Stoddard, Favour A. Oladimeji, Virginia Spanoudaki, Kasturi Chakraborty, Savan K. Patel, Heather E. Fleming, Christopher S. Chen, Sangeeta N. Bhatia

## Abstract

Liver transplantation remains the standard of care for end-stage liver failure, yet it is constrained by donor scarcity, surgical complexity, and restricted access for many patients. Cell-based therapies offer a potential alternative, yet their translation has been hindered by low engraftment, poor localization, and a lack of delivery strategies that are both effective and minimally invasive. To address these challenges, we developed a new approach termed **INSITE** (**<u>In</u>**jected **<u>S</u>**elf-assembled **<u>I</u>**mage-guided **<u>T</u>**issue **<u>E</u>**nsembles), an injectable platform composed of primary human hepatocytes and hydrogel microspheres that can be delivered by image-guided injection and assembled in situ into supportive, vascularizable scaffolds. In vivo, ultrasound-guided delivery into an ectopic site enabled precise graft localization, persistent visibility under noninvasive imaging, and vascular integration. Hepatocytes within these niches remained confined to the scaffold and maintained long-term functional activity. Furthermore, tuning material properties allowed control over scaffold remodeling and vascular recruitment, providing a means to enhance graft function. By integrating image-guided delivery with a modular and supportive scaffold, INSITE establishes a clinically compatible strategy for advancing minimally invasive cell therapies.

## Introduction

Organ transplantation is often the only curative solution for end-stage organ failure, yet it is accessible to only a small fraction of patients who need it. For several vital organs such as the liver, kidney, and pancreas, donor supply falls far short of demand, and many patients die while waiting for a suitable match^1,2^. Even when a donor organ is available, the procedure itself poses major barriers: transplantation requires highly specialized surgical teams and infrastructure, advanced perioperative care, and patients stable enough to tolerate major surgery. These challenges underscore an urgent need for scalable, less invasive alternatives that could expand access to tissue replacement therapies. Towards this, cell therapies and tissue engineering represent attractive opportunities, offering the possibility to restore function without the limitations of donor availability^3,4^. Minimally invasive delivery strategies would further accelerate this transition, creating opportunities to treat patients who would otherwise never benefit from transplant surgery.

To overcome the limitations of surgical implantation, researchers have explored a variety of strategies for building therapeutic tissues directly within the body. In some cases, cells have been injected directly, sometimes intravascularly, but cell survival and engraftment are typically poor without a defined niche^5^. To improve engraftment, biomaterial approaches have been developed; however, most conventional scaffold-based strategies require invasive implantation^6^. This limitation has been circumvented by the development of injectable systems, such as hydrogels that encapsulate cells, aided by photopolymerization or in situ bioprinting, to provide supportive scaffolding at the target site^7–9^. While these methods have demonstrated the potential of minimally invasive tissue engineering, each faces trade-offs, including limited host integration, poor spatial control, and technical difficulties in applying these strategies within deep-seated tissues. Moreover, without image-guided delivery, these approaches lack precision and efficiency, and do not provide control over the niche into which cells and materials localize, nor how effectively they integrate with host tissue. Granular hydrogels offer a promising solution to many of these challenges. Composed of microscale building blocks, in that they can be manipulated as a fluid via shear-induced flow, and once the shear is removed the microparticles jam together, forming into a stable, microporous scaffold that assembles in situ^10,11^. These scaffolds provide mechanical support while enabling nutrient exchange and cell recruitment, thus accelerating integration with host^12^. Until now, granular hydrogels have been largely applied in acellular contexts in which porosity can aid wound healing and tissue repair by permitting host cell migration, such as to promote cellular infiltration in skin wounds^13,14^, to support neural progenitor cell migration into the cavities of brain lesions^15^, to facilitate remodeling after myocardial infarction^16^, or to serve as therapeutic molecule carriers during bone repair^17^. Using granular hydrogels to engineer a functional transplant niche for minimally-invasive delivery of therapeutic cells could significantly advance solid organ cell therapies.

Here, we introduce INSITE, an injectable, *in situ*-assembling niche using gelatin methacryloyl (GelMA) microspheres designed to support hepatocyte transplantation. We demonstrate that primary human hepatocytes (PHHs) can be incorporated into these niches and delivered using ultrasound-guided injection. Image-guided injection allows the grafts to be precisely localized, overcoming the dispersion and inefficiency associated with traditional cell delivery approaches. Following injection, the niches remain confined within the target site, recruit vasculature from the host, can be imaged non-invasively, and maintain long-term hepatocyte function. By constructing defined niches that condition the local microenvironment, this approach makes target tissues more receptive to engraftment and long-term function. Importantly, this strategy transforms liver tissue engineering from a procedure that traditionally requires open surgery to one that could be performed minimally-invasively, using widely available clinical tools. More broadly, this platform highlights how injectable, self-assembling materials can help shift cell-based therapies from surgery-dependent interventions to scalable treatments for organ failure.

## Results

### GelMA microspheres assemble into injectable, macroporous scaffolds compatible with ultrasound-guided in vivo delivery

Towards our goal of developing in situ assembling, injectable liver grafts, we fabricated granular hydrogels composed of GelMA microspheres. GelMA prepolymer solution was emulsified as aqueous droplets in a continuous oil phase using a step-emulsification microfluidic device (**Supplementary Fig. 1A, Supplementary Video 1**), and the resulting droplets were subsequently crosslinked by UV exposure to form stable GelMA microspheres that distributed around a mean diameter of 110 μm (**Fig. 1A**).

**Figure 1.**
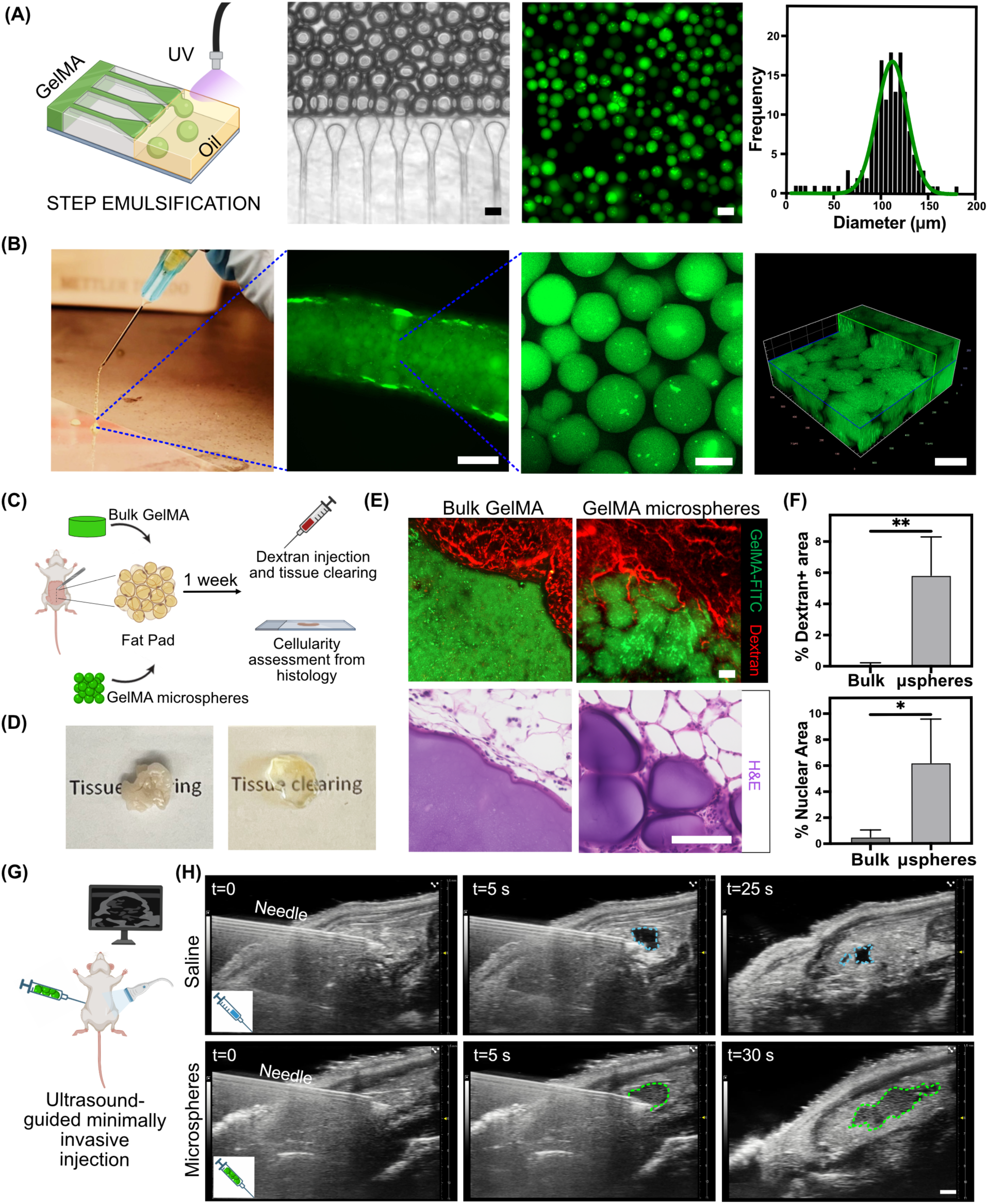
GelMA microspheres form injectable, macroporous scaffolds that integrate with host tissue and can be delivered under ultrasound guidance. (A) Step-emulsification microfluidics for fabrication of GelMA microspheres and their characterization. Aqueous phase hydrogel precursor is emulsified in oil phase into droplets that are crosslinked via UV exposure to yield the GelMA microspheres (scale bars 100 µm). The microspheres have an average diameter of 110 μm and Coefficient of Variation of 22.8% (B) Extrusion of microspheres packed into a slurry through a 25G needle (far left). Confocal imaging of the extruded slurry (left middle) demonstrating preservation of interstitial voids post-injection (middle right), and 3D reconstruction (far right) (scale bar 500 µm for the left middle image, 100 µm for others). (C) Schematic of surgical implantation of bulk GelMA hydrogel or GelMA microspheres into the fat pad of NSG mice, followed by dextran perfusion, tissue clearing, and histological analyses one week later. (D) Representative macroscopic images of explanted tissue with implanted hydrogels before and after clearing. (E) Cleared whole mounts showing FITC-labeled GelMA (green) and perfused dextran (red), and H&E sections of bulk GelMA and microsphere scaffolds (scale bars 100 µm). (F) Quantification of dextran-positive vessel area fraction (top) and nuclear infiltration (bottom) Multiple fields were stitched to generate a mosaic image of the graft, and the stitched image was analyzed to yield one quantitative data point per animal by calculating Texas Red^+^ and Hoechst^+^ area, respectively within the graft area. Data are mean ± SD; unpaired two-tailed Student’s t-test (n = 4 animals per group) (G) Schematic of ultrasound-guided injection of GelMA microspheres into the fat pad. (H) Representative B-mode ultrasound images showing needle placement (t=0), bolus appearance (t=5 s, dashed blue outline for saline and green outline for GelMA microspheres), and signal after needle removal (scale bar 1 mm).

To evaluate injectability, GelMA microspheres were first centrifuged to form a densely packed slurry, which was then loaded into a syringe and extruded through a 25-gauge needle (**Fig. 1B**). The packed microspheres were readily extruded through the needle, confirming their injectability. Confocal imaging showed that interstitial voids between microspheres were preserved (**Fig. 1B)**, and quantitative analysis confirmed that the void fraction remained unchanged before and after injection (**Supplementary Fig. 1B**).

We next examined the viscoelastic properties of packed GelMA microspheres. At low strain, the storage modulus exceeded the loss modulus, consistent with solid-like properties of a jammed microsphere network (**Supplementary Fig. 1C**). When subjected to higher strain, the material underwent yielding, marked by a sharp decrease in storage modulus, indicating a transition toward a fluid-like state (**Supplementary Fig. 1C**). Frequency sweep demonstrated that the storage modulus remained higher than the loss modulus across a broad range, confirming predominantly elastic behavior under oscillatory loading (**Supplementary Fig. 1D**). Flow sweep analysis further revealed shear-thinning, with viscosity decreasing as shear rate increased (**Supplementary Fig. 1E**). Together, these measurements establish that packed microspheres behaved as elastic solids at rest but undergo shear-thinning under deformation, enabling injection through a needle while re-forming into a stable, porous scaffold once stress is removed.

We next assessed whether acellular granular GelMA hydrogels supported host cell integration following implantation in vivo. To this end, either monolithic bulk GelMA hydrogels or granular hydrogels composed of densely packed microspheres were implanted into the perigonadal fat pad of NSG mice (**Fig. 1C**). We selected this site because our prior studies have shown that hepatocytes survive better in intraperitoneal fat tissues compared to subcutaneous locations, making it a relevant site for testing hepatocyte graft niches^18,19^. One week after implantation, mice were injected intravenously with fluorescent dextran to visualize perfused vasculature. Tissue clearing (**Fig. 1D**) followed by whole-mount imaging revealed that bulk gels maintained a distinct interface with host tissue and were largely avascular, whereas packed microspheres exhibited vascular infiltration into the interstitial voids (**Fig. 1E**). Histological analysis confirmed increased cellularity in packed microspheres (**Fig. 1E**). Quantitative analysis showed a significantly higher vessel area fraction and nuclear infiltration in microspheres-based graft compared to bulk controls (**Fig. 1F**).

As a proof-of-concept for image-guided delivery, we tested whether GelMA microspheres could be administered under ultrasound guidance. Packed microspheres were injected into the perigonadal fat pad under real-time ultrasound visualization (**Fig. 1G**). The hydrogel suspension appeared as a moderately echogenic bolus, distinguishable in contrast from the surrounding fat, allowing accurate placement within the target tissue (**Fig. 1H**). The injected scaffolds remained detectable by ultrasound for several weeks (**Supplementary Fig. 2A**), with visibility likely arising from acoustic scattering from the numerous interfaces formed within the heterogeneous microsphere scaffold (endogenous contrast). This property underlies the ability of the INSITE platform to remain trackable over time under noninvasive imaging. Furthermore, we employed nonlinear contrast-enhanced ultrasound with non-targeted microbubble contrast agents. The microbubbles were administered intravenously to visualize host circulation, and perfusion signals were detected within the graft region consistent with vascular connectivity of the implanted granular hydrogels (**Supplementary Fig. 2B, Supplementary Video 2**). Taken together, these findings demonstrate that GelMA microspheres can be formulated into injectable, macroporous scaffolds that promote vascular and cellular integration in vivo and can be precisely administered via ultrasound-guided delivery, positioning this strategy as a clinically compatible platform for creating injectable hepatocyte transplant niches. Beyond their accurate placement, the macroporous architecture facilitates vascular recruitment, effectively creating a niche that renders host tissue more hospitable for transplanted cells. Building on this foundation, we next investigated whether these injectable scaffolds could provide a supportive microenvironment for human hepatocytes.

### GelMA microspheres provide a supportive scaffold for hepatocyte–fibroblast aggregates in vitro

To investigate whether packed GelMA microspheres could serve as a supportive matrix for hepatocyte culture, we aggregated primary human hepatocytes (PHHs) with neonatal human dermal fibroblasts (NHDFs) for three days prior to encapsulation (**Fig. 2A**). Our previous studies have shown that juxtacrine signaling from stromal cells like fibroblasts helps stabilize the phenotype and function of primary hepatocytes^20–22^. Aggregates were then collected and combined with GelMA microspheres to generate a composite slurry that was cultured in vitro for up to 2 weeks.

**Figure 2.**
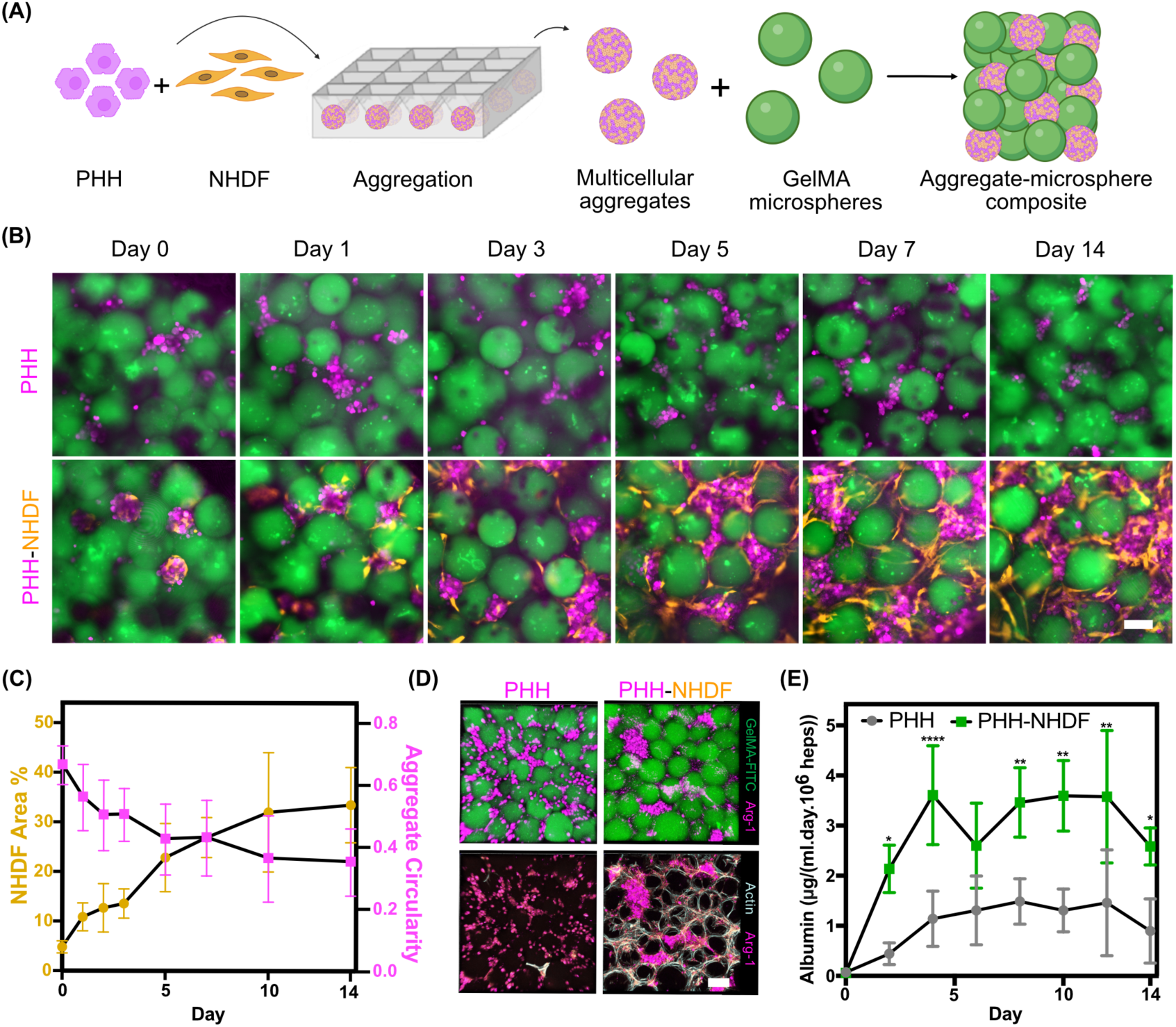
Injectable GelMA microspheres support the culture and functional maintenance of primary human hepatocytes in vitro. (A) Schematic of the strategy to incorporate Primary Human Hepatocytes (PHHs) into a slurry of GelMA microspheres to generate engineered grafts. PHHs were aggregated with neonatal human dermal fibroblasts (NHDFs) for 3 days in round-bottom wells to yield multicellular aggregates that, once mixed with densely-packed GelMA microspheres, yield an aggregate-microsphere composite slurry that can be cultured in vitro. (B) Longitudinal fluorescence imaging of PHH-only and PHH-NHDF aggregates among microspheres over the course of 14 days (scale bar 100 µm). (C) Quantification of NHDF area fraction (RFP^+^ area in 8-17 fields of interest (FOI) per time point) and aggregate circularity (4*pi*Area/Perimeter^2^ of aggregates in 8-17 fields of interest (FOI) per time point) over time in PHH-NHDF co-cultures. (D) Immunofluorescence at day 14 showing Arg-1^+^ (magenta) hepatocytes along with F-actin to mark other cell populations within the culture (scale bar 100 µm). (E) Albumin secretion from PHH-only vs PHH-NHDF aggregates cultured in GelMA microspheres. Data are mean ± SD; two-way repeated measures ANOVA with Šidák’s post hoc test.

Longitudinal fluorescence imaging revealed several differences between PHH-only and PHH-NHDF aggregates cultured in GelMA microspheres (**Fig. 2B**). PHH aggregates cultured without fibroblasts failed to maintain spheroid integrity, fragmenting into small clusters and single cells with limited remodeling within the microsphere scaffold. By contrast, in co-cultures, fibroblasts migrated outward from the aggregates, adopted an elongated morphology, and spread across the surfaces of surrounding microspheres. This resulted in the establishment of a stromal network that bridged adjacent aggregates and microspheres. In parallel, hepatocytes remained clustered within spheroids, but the spheroids themselves progressively conformed to the interstitial voids between microspheres, resulting in a gradual reduction in circularity over the culture period. Quantitative analysis confirmed a decrease in aggregate circularity in co-cultures and a progressive increase in fibroblast coverage within the scaffold (**Fig. 2C**).

To further characterize these cultures, aggregates were analyzed by immunofluorescence after 14 days of culture. Hepatocytes expressed the mature hepatocyte marker arginase-1 (Arg-1), while fibroblasts were distributed throughout the scaffold in close association with hepatocyte clusters (**Fig. 2D**). In PHH-only cultures, a very small number of mesenchymal-like cells were observed, attributable to the contamination of non-parenchymal cells (NPCs) during PHH isolation. However, these cells did not organize into supportive stromal networks as seen in co-cultures. Analysis of secreted albumin in the culture media revealed significantly higher levels in PHH–NHDF aggregates compared to PHH-only controls (**Fig. 2E**). PHHs cultured with or without fibroblasts exhibited comparable urea production, as quantified by urea assays of conditioned media at Day 14 (49.34 ± 25.53 µg/mL/day/10⁶ cells for hepatocytes alone versus 46.57 ± 11.57 µg/mL/day/10⁶ cells for hepatocytes co-cultured with fibroblasts). PHHs in both conditions also maintained sustained drug-metabolizing capacity, as assessed by CYP3A4 activity following rifampicin induction. While CYP3A4 activity was detectable in both groups, levels were higher in PHHs co-cultured with fibroblasts (**Supplementary Fig. 3A**), and this difference increased over time in culture, consistent with previous studies that concluded that PHHs depend on fibroblast interactions to maintain functional output^23–25^. In addition, hepatocytes retained transporter-mediated functional activity, as demonstrated by indocyanine green uptake, which was observed selectively in hepatocytes but not in fibroblasts (**Supplementary Fig. 3B**). Collectively, these results demonstrate that GelMA microspheres provide a supportive microenvironment that stabilizes hepatocyte–fibroblast aggregates and preserves hepatocyte function in vitro, highlighting their potential for establishing transplantable niches in vivo.

### Engineering an injectable hepatocyte transplant niche enables efficient engraftment and long-term maintenance of ectopically transplanted hepatocytes

To test the ability of GelMA microspheres to establish a transplant niche for long-term hepatocyte engraftment and function, we compared three conditions in vivo: PHH-only aggregates, PHH–NHDF aggregates, and PHH–NHDF aggregates delivered with GelMA microspheres. Aggregates were suspended in equal volumes of either media (PHH-only and PHH–NHDF) or microsphere slurry (PHH–NHDF) immediately before injection. Each formulation was injected into the perigonadal fat pad of NSG mice, and grafts were monitored for 8 weeks using ultrasound imaging before endpoint analysis of hepatocyte survival, vascularization, and integration with host tissue (**Fig. 3A**).

**Figure 3.**
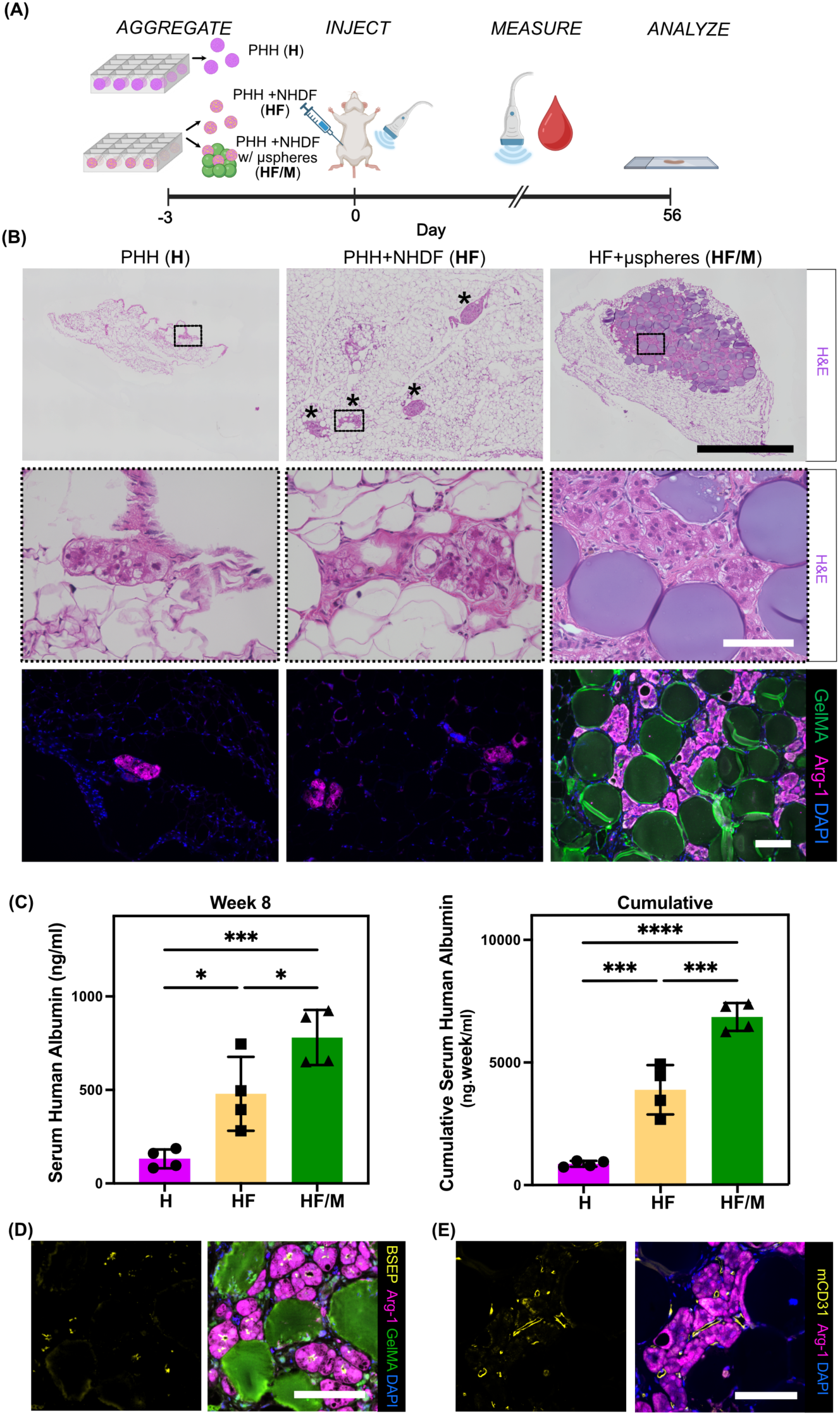
Injectable GelMA microspheres enable efficient and long-term engraftment of primary human hepatocytes in vivo. (A) Schematic of experimental design comparing PHH aggregates (H), PHH-NHDF aggregates (HF), and PHH-NHDF aggregates with GelMA microspheres (HF/M) in vivo. Aggregates were generated 3 days prior to injection, followed by delivery into the fat pad, longitudinal monitoring via ultrasound and blood draws, and endpoint analyses at 8 weeks. (B) Representative H&E sections (scale bars, top, 1 mm; dotted-outlined insets (middle), 100 µm) showing limited hepatocyte clusters in H, dispersed aggregates in HF (asterisks), and compact tissue-like grafts in HF/M, and immunofluorescence (bottom) for hepatocyte marker Arg-1^+^ (magenta) hepatocytes (scale bar 100 µm). (C) Serum human albumin levels at week 8 (left) and cumulative output over 8 weeks based on weekly blood samples (right). Data are mean ± SD; one-way ANOVA with Tukey’s post hoc test. (D) Immunofluorescence showing BSEP (yellow) canalicular localization in Arg-1^+^ (magenta) hepatocytes within HF/M grafts (scale bar 100 µm). (E) Immunostaining demonstrating close association of Arg-1^+^ (magenta) hepatocytes with mCD31^+^ (yellow) host vasculature (scale bar 100 µm).

During injection, distinct differences were observed between groups when visualized under ultrasound. As with acellular microspheres, grafts containing GelMA microspheres appeared as moderately echogenic structures confined to the fat pad, forming a localized deposit that could be readily visualized (**Supplementary Fig. 4A, Supplementary Video 3**). By contrast, both PHH-only and PHH–NHDF aggregates suspended in media behaved like a hypoechoic fluid bolus, dispersing through the adipose tissue without forming a stable structure (**Supplementary Fig. 4A, Supplementary Video 4, 5**).

Longitudinal imaging using ultrasound and quantification of the relative contrast observed revealed that only microsphere-containing grafts remained visible beyond the day of injection, which enabled the quantification and longitudinal tracking of the graft volume (**Supplementary Fig. 4B,C**). In the PHH-only and PHH–NHDF conditions, the injected material could not be detected aside from the transient hypoechoic signal during injection. By contrast, microsphere grafts were detectable as localized structure for the full 8-week study period (**Supplementary Fig. 4B,C**). Nonlinear contrast-enhanced ultrasound with non-targeted microbubble contrast agents demonstrated perfusion signals within the graft, further supporting vascular connectivity (**Supplementary Fig. 4D**). Thus, ultrasound imaging not only confirmed persistence of microsphere grafts but also provided a means of tracking vascular integration.

End-point histological analyses corroborated these ultrasound observations (**Fig. 3B**). In both PHH-only and PHH–NHDF aggregate groups, only a few scattered clusters of hepatocytes could be identified within the fat pad. In contrast, aggregates delivered with microspheres formed compact, tissue-like niches that retained structure and localized the hepatocytes. Analysis of serum albumin output revealed that the levels of circulating human albumin for PHH–NHDF aggregates with microspheres consistently remained higher than in other cohorts (**Supplementary Fig. 5A**), and the cumulative secretion (area under the curve) was highest in mice receiving PHH–NHDF aggregates with microspheres, significantly exceeding both PHH-only and PHH–NHDF aggregates (**Fig. 3C**). At week 8, albumin levels remained significantly elevated in the microsphere group compared to aggregates delivered without microspheres (**Fig. 3C**).

Immunostaining confirmed expression of the hepatocyte marker Arg-1 within the grafts (**Fig. 3B**). Hepatocytes also exhibited canalicular bile salt export pump (BSEP) localization, consistent with preservation of mature lineage and polarity (**Fig. 3D**); this was also observed in the PHH–NHDF grafts (HF) but not in the grafts of the PHH-only (H) cohort (**Supplementary Fig. 5B**). Engrafted hepatocytes were found in close association with CD31⁺ host vasculature in every condition examined, indicating integration with the host circulation (**Fig. 3E, Supplementary Fig. 5C**). Notably, despite these shared indicators of hepatocyte polarity and vascular integration, the predominant distinction between treatment groups was the spatial organization of the engrafted cells: in the condition with microspheres, hepatocytes were localized within the scaffold regions defined by the microspheres, whereas in the other conditions hepatocytes were present only as scattered clusters, suggesting limited and uncontrolled engraftment. Further characterization of the PHHs within the PHH–NHDF aggregates with microspheres conditions revealed that the cells expressed CYP3A4 as validated through immunostaining (**Supplementary Fig. 5D**). TUNEL (Terminal deoxynucleotidyl transferase dUTP nick end labeling) staining performed on explants showed minimal cell death (< 1.2% TUNEL+ within the graft), and co-staining for TUNEL with Arg-1 showed no overlap between the detected TUNEL signal and hepatocytes (**Supplementary Fig. 5E**). Overall, these findings demonstrate that the INSITE platform establishes a stable, vascularized, and functionally organized hepatocyte niche that can be delivered minimally invasively and monitored noninvasively over time, underscoring its potential as a clinically relevant platform for hepatocyte transplantation.

### Tuning degradability of microspheres allows control over scaffold persistence and graft function

Engineered scaffold remodeling by the host is crucial for regulating graft integration, vascularization and function. Towards this, we investigated whether the degradability of GelMA microspheres could be tuned to influence these outcomes. Microspheres were fabricated with different degrees of GelMA crosslinking to generate formulations with high- and low-degradability (**Fig. 4A**). Enzymatic degradation assays revealed that the high-degradability formulation underwent more rapid breakdown when incubated with collagenase in vitro, whereas the low-degradability formulation (used in the experiments described so far) underwent significantly slower degradation over a period of 6 hours (**Supplementary Fig. 6A**).

**Figure 4.**
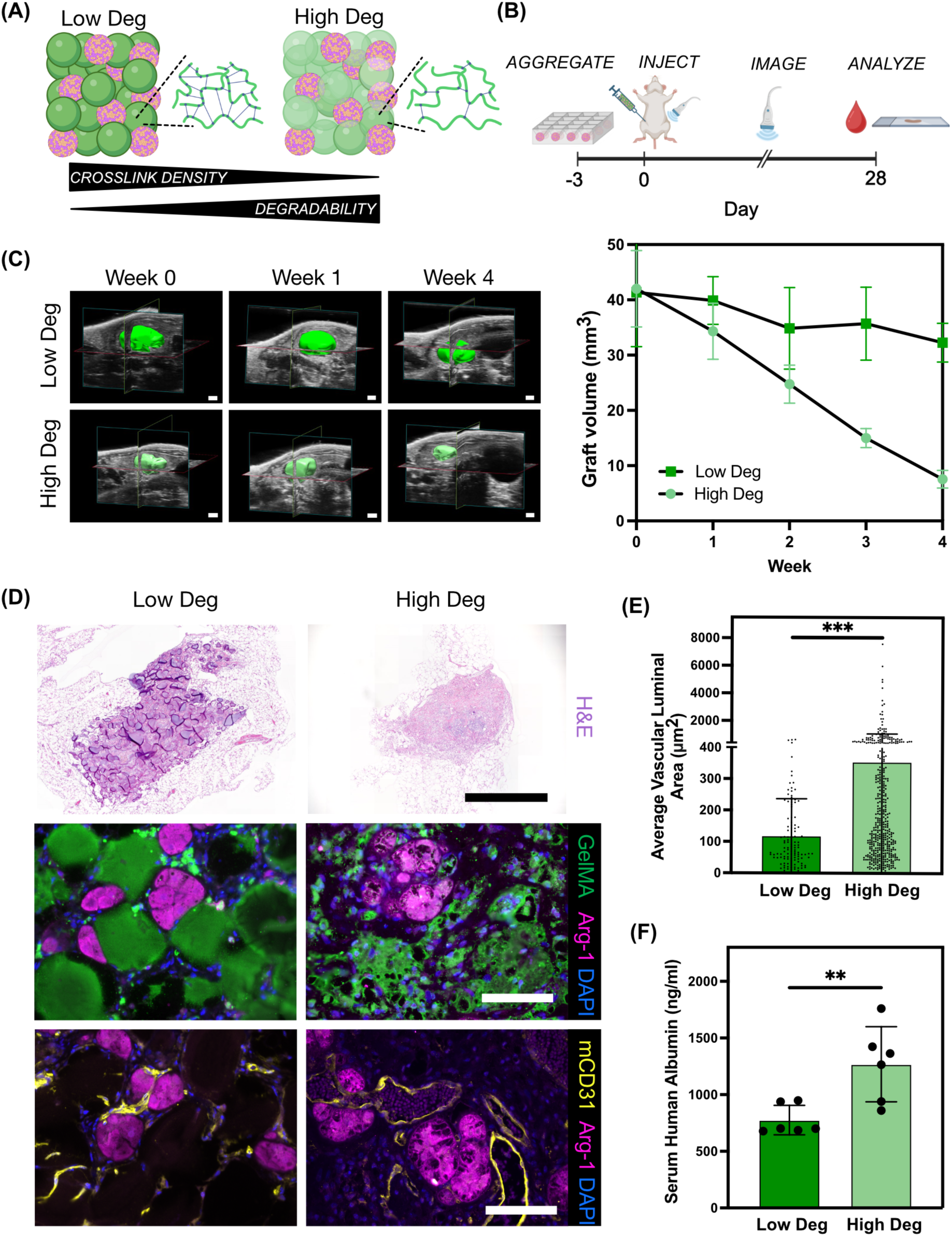
Scaffold degradability influences graft remodeling, vascularization, and hepatocyte function. (A) Schematic of GelMA microspheres with varied degradability based on either higher crosslink density (Low Deg) versus lower crosslink density (High Deg). (B) Experimental timeline showing aggregate preparation, ultrasound-guided injection, longitudinal imaging, and analyses at day 28. (C) Representative 3D ultrasound reconstructions of grafts and longitudinal quantification of microsphere volume over time (scale bars 1 mm). Data are mean ± SD (n = 6 animals per group pooled from 2 independent experiments with 3 animals each). (D) Representative H&E images (top panel), immunofluorescence showing Arg-1^+^ (magenta) hepatocytes retained within GelMA (green) scaffolds (middle panel), and mCD31^+^ (yellow) vasculature associated with Arg-1^+^ (magenta) hepatocytes in Low Deg and High Deg niches (bottom panel) (scale bars, top panel 1 mm, middle and bottom panel 100 µm). (E) Quantification of average vascular luminal area (calculated by tracing CD31^+^ areas from >10 fields from 3 mice in each group). Data are mean ± SD; unpaired two-tailed Student’s t-test. (F) Serum human albumin levels at day 28. Data are mean ± SD (n=6 mice pooled from 2 independent experiments with 3 mice each); unpaired two-tailed Student’s t-test.

Both variants remained injectable and when delivered via ultrasound-guided injection with PHH–NHDF aggregates, remained localized and echogenic within the perigonadal fat pad post-injection (**Fig. 4B**). Longitudinal ultrasound monitoring demonstrated that high-degradability gels resorbed rapidly, with significant loss of detectable volume by four weeks, while low-degradability gels only slightly lost volume during the same period (**Fig. 4C**). Histological analysis showed that low-degradability microspheres maintained structural integrity and persisted in vivo, whereas high-degradability gels were extensively remodeled while still retaining hepatocytes within the niche (**Fig. 4D**). In high-degradability gels, vessels exhibited strikingly larger average luminal cross-sectional area per vessel and extended beyond the original void spaces between microspheres into the remodeled niche, whereas in low-degradability gels vessels remained confined to the interstitial voids (**Fig. 4D, E).** In parallel, F4/80 staining revealed that macrophages in high-degradability conditions dispersed more widely across the graft, whereas in low-degradability gels their presence was largely restricted to the void spaces between microspheres (**Supplementary Fig. 6B**). To study macrophage phenotypes, we stained for CD86 and CD206 expression to assay for the presence of pro-inflammatory and pro-regenerative populations, respectively. We found that macrophages in Low Deg conditions showed a trend towards a more pro-inflammatory macrophage phenotype compared to High Deg (% CD86⁺ area/graft), whereas differences in pro-regenerative macrophages (% CD206⁺ area/graft) were comparatively modest, but trended higher in the High Deg condition (**Supplementary Fig. 6C**).

To determine whether scaffold degradability impacted hepatocyte function, serum albumin levels were measured in mice transplanted with PHH–NHDF aggregates delivered in either high- or low-degradability gels. At week 4, higher albumin secretion was observed in the high-degradability group (**Fig. 4F**) suggesting improved graft function. These results suggest that scaffold degradation kinetics can be tuned to balance persistence and remodeling, and that these dynamics can be tracked noninvasively with ultrasound imaging. Together, these findings add an important dimension to the utility of GelMA microspheres, demonstrating that degradability is a tunable parameter that influences the function of transplanted hepatocytes.

## Discussion

This study establishes INSITE, a hydrogel microspheres-based, minimally-invasive, image-guided platform to engineer niches that support the function of transplanted human hepatocytes. Using ultrasound-guided delivery, we show that microspheres can be injected into ectopic sites, remain localized, and provide inherent echogenic contrast that enables longitudinal monitoring without additional labels. Within these niches, hepatocytes survived, retained identity, and contributed to stable, tissue-like structures over long term. Further, adjustments of microsphere degradability altered scaffold remodeling and vascular integration, demonstrating that material properties can serve as tunable levers to modulate graft performance.

The clinically-compatible strategies demonstrated in this study offer a possible route to overcome long-standing barriers in hepatocyte transplantation. There is an urgent need for such advances because current options for liver replacement remain limited^26^. Transplantation is the only curative therapy for end-stage liver failure, but access is restricted by organ shortages, surgical complexity, and the need for specialized centers. Cell-based therapies offer an attractive alternative, yet clinical experience with hepatocyte infusion highlights persistent limitations of cell dispersion, inefficient engraftment, and failure to sustain long-term function^27^. Attempts to engineer replacement liver tissue have sought to improve outcomes by pre-assembling tissue-engineered grafts ex vivo^28–32^, but such constructs often require open surgical implantation, limiting feasibility and translation. Other biomaterial strategies have aimed to build grafts directly in vivo, including photopolymerizable bulk gels^33^, sprayable bioinks^34^, and in vivo bioprinting approaches^35,36^. While many injectable hydrogels and in situ crosslinking systems have shown promise, they often rely on rapid post-delivery gelation, which can complicate injectability, limit spatial precision, and delay early host integration due to the lack of macroporosity at implantation^37^. Photocrosslinking, one of the most common in situ strategies, is fundamentally constrained in deep tissues by light attenuation: ultraviolet light typically penetrates only a few millimeters, while longer-wavelength visible and near-infrared light provides only modest improvements without invasive delivery methods such as fiber-optic or endoscopic illumination^38,39^. X-ray–based crosslinking has recently been shown to enable polymerization at tissue depths of up to ∼2 cm, although safety, dosing, and long-term biological effects of ionizing radiation remain to be established in therapeutic settings^40^.

In vivo bioprinting approaches can provide architectural control but generally require specialized equipment and face limitations in deep tissues. Extrusion-based in vivo bioprinting is largely restricted to superficial or surgically-exposed sites, while stimulus-responsive bioprinting strategies, including light-based systems, share similar depth constraints^35^. Sound-based in vivo printing has recently demonstrated material deposition within deeper tissues, including the bladder, but remains technically complex and infrastructure-intensive^36^. By contrast, INSITE relies on pre-formed hydrogels and can be delivered to deep tissue sites accessible by ultrasound-guided injection in clinical settings, assembling without reliance on post-delivery gelation.

Injectable granular hydrogels have emerged as promising materials for tissue repair, with different formulations tested in sites including the skin^13,41^, brain^15,42^, heart^16,43^, cartilage^44,45^, bone^17,46^ etc. Composed of microgels as building blocks, their macroporous architecture has been shown to accelerate repair by enabling host cell infiltration and migration without requiring scaffold degradation, distinguishing them from bulk hydrogels that integrate only after remodeling while risking encapsulation and limited host integration^12,47–50^. These hydrogels have also been used as carriers for delivering therapeutic molecules, and more recently, combined with chondrocyte spheroids for cartilage tissue engineering in vitro and ex vivo^44^. Building upon these advances, the present strategy establishes granular hydrogels composed of GelMA microspheres as minimally invasive, image-guided niches for therapeutic cell transplantation in vivo.

These findings raise the possibility that hepatocyte transplantation could be pursued without reliance on major surgical procedures. Instead, the ability to deliver and monitor grafts under real-time ultrasound guidance suggests that such therapies could be performed by interventional radiologists using minimally-invasive techniques already in routine clinical practice. Unlike organ transplantation, which requires specialized surgical teams, intensive perioperative care, and centralized transplant centers often far from patients, ultrasound-guided, injectable grafts could be implemented in a wider range of clinical settings with reduced infrastructure demands. Moreover, by creating a physically-defined niche, this strategy addresses one of the major limitations of current clinical hepatocyte infusions, during which injected cells disperse and fail to engraft efficiently. The lower procedural burden of using injectable hydrogel microspheres, coupled with improved localization and engraftment, would not only decrease cost but also broaden patient eligibility, particularly for those unable to tolerate major surgery. Overall, shifting from open surgery to image-guided injection has the potential to significantly expand access to cell-based liver therapies.

A limitation of the current approach is that the niche visualization by ultrasound relies on the endogenous contrast generated by multiple interfaces within the microsphere scaffold. While this physical trait enables monitoring of graft localization and persistence, it does not allow discrimination between the microspheres and the hepatocyte aggregates interspersed between them. Ideally, future iterations of this platform could incorporate microspheres loaded with defined contrast agents to visualize the scaffold, and paired with orthogonally-labeled hepatocytes and/or supporting cell types, thereby enabling independent tracking of both the niche and the transplanted cells.

Future studies will include an exploration of the maximum size of the niche that can be generated with this approach, as scaling to larger volumes may introduce additional considerations related to diffusion, vascularization, and structural stability, as well as injection sites. A parallel challenge in scale-up is the sourcing of sufficient numbers of functional cells^51^. While iPSC- and other stem-cell–derived hepatocyte-like cells have shown promise, they generally fail to achieve fully mature hepatocyte identity^52^. Recent advances in expanding primary human hepatocytes in 2D, as organoids, and in vivo suggest that proliferative PHHs could provide a viable cell source for generating clinically-relevant graft sizes^53–57^. The injectable, modular, and image-guided nature of INSITE would enable dose scaling either by increasing the injected volume, or by distributing cells across multiple discrete niches - potentially avoiding the need for a single large graft that could be vulnerable to the development of diffusion-limited necrotic cores. Importantly, the minimally-invasive delivery paradigm also raises the possibility of serial dosing, in which additional niches could be administered over time based on functional circulatory biomarkers. This capacity would allow dosing to be adapted in response to graft performance, and would otherwise be difficult to implement in settings that rely on surgical implantations. Future iterations of the platform could further improve engraftment and enable greater implant volumes by incorporating endothelial cells directly within the graft to accelerate vascularization. With respect to delivery, ultrasound-guided injection can be adapted for scaling graft size by withdrawing the needle at a controlled speed while modulating the extrusion rate to generate elongated, high–aspect-ratio grafts that reduce diffusion distances. That said, this approach is necessarily constrained in vivo by the limited freedom to move the needle in multiple directions within tissue. Nonetheless, ultrasound guidance can enable the placement of multiple grafts at defined size limits within the target tissue, and real-time visualization can ensure that adjacent grafts remain spatially distinct rather than merging into a single diffusion-limited mass. The ratio of microspheres to hepatocyte aggregates could also be further tuned to optimize graft architecture and function. Adjusting this balance may allow control over the interplay between structural support, vascular infiltration, and cell density, providing a means to tailor the niche for desired therapeutic outcomes. While the present findings suggest that degradability enhances graft function, the mechanisms underlying this effect remain to be fully elucidated. It is possible that differences in vascular remodeling and immune infiltration contribute to the observed outcomes. Other recent studies have demonstrated that the rate and mechanism of microgel degradation can influence cell spreading, proliferation, extracellular matrix deposition, and macrophage polarization^58–60^. The increased persistence of the higher crosslinked GelMA microspheres in the Low Deg condition may promote sustained macrophage–microsphere interactions, potentially leading to prolonged retention of pro-inflammatory macrophages at the site. This interpretation is consistent with prior reports showing that stiffer GelMA hydrogels can bias macrophages toward an M1 phenotype^61,62^, suggesting that crosslinking density and the resulting material persistence in vivo could be an important contributing factor to the observed polarization trends, even within the constrained immune environment of this model. A decrease in pro-inflammatory macrophages in the High Deg graft would increase the relative proportion of pro-regenerative macrophages, which likely promotes enhanced vascularization, as evidenced by the presence of larger vessels observed within the High Deg grafts. Improved vascularization may in turn support better hepatocyte function in the High Deg graft. However, these results should be interpreted with caution, as immunodeficient mouse models do not fully recapitulate the complexity of macrophage polarization observed in intact immune systems, and definitive conclusions will require validation in immunocompetent models. Furthermore, differences in macrophage polarization can be modest when assessed using classical pro-inflammatory and pro-regenerative markers, reflecting the fact that macrophage activation occurs along a continuum rather than as discrete states^63^. As demonstrated in multiple studies, in vivo macrophage phenotypes frequently do not conform to strict M1/M2 classifications, but instead acquire dynamic, context-dependent intermediate states with studies showing that expression of CD86 and CD206 are not mutually exclusive^64^. In addition to macrophage polarization, elements of the foreign body response are expected to be reduced here relative to immunocompetent hosts, potentially affecting scaffold remodeling and vascularization. Further studies would be needed to define how macrophage polarization and the foreign body response influence scaffold integration, vascularization, and long-term hepatocyte function in immunocompetent or humanized immune models.

Looking forward, the modularity of this system could be leveraged to incorporate microspheres of varying degradability within the same graft, where highly degradable components promote remodeling and vessel ingrowth, while more stable counterparts provide long-term mechanical support. Additional opportunities include incorporating other relevant cell types beyond fibroblasts, which are included here to stabilize hepatocyte phenotype and function through juxtacrine signaling. Extensive prior work has demonstrated that fibroblast coculture is sufficient to sustain long-term primary human hepatocyte function across 2D systems and 3D formats, both in vitro and in vivo^18,20–22,65^. Together with fibroblast-mediated support, the granular hydrogel scaffold contributes architectural advantages in vivo accelerating vascularization. Building on this framework, future studies could incorporate additional cell types such as biliary epithelium^66^, and guide their organization within interstitial voids to recreate aspects of the native hepatic biliary tree. Additionally, the microspheres themselves could be adapted for use as bio-inks in 3D bioprinting^67,68^, enabling the fabrication of constructs that more closely recapitulate the hierarchical architecture of the liver.

Finally, INSITE could be extended to other cell therapies that face similar barriers to engraftment/whole organ transplantation. For example, pancreatic islet transplantation for type 1 diabetes is currently performed by intravascular infusion into the liver. However, this practice also suffers from many of the same limitations as hepatocyte delivery, including cell dispersion, poor engraftment, and progressive loss of function^69^. By replacing uncontrolled intravascular dispersal with an image-guided, injectable niche, the same principles demonstrated here for hepatocytes could in the future provide a supportive, vascularized environment for islets, potentially improving localization, survival, and long-term function.

In summary, this work establishes hydrogel microspheres as a minimally invasive, image-guided platform for engineering functional hepatocyte transplant niches. By replacing open surgery with interventional delivery, supporting long-term graft function, and enabling tunable vascular integration, this approach addresses key barriers that have limited a wider adoption of hepatocyte transplantation. More broadly, injectable, self-assembling niches represent a significant step toward regenerative treatments that are more accessible, scalable, and available to patients who may not receive a donor organ.

## Materials and methods

### GelMA Synthesis

GelMA was synthesized by reacting porcine type A gelatin (Millipore Sigma, 1890) with methacrylic anhydride (MAA) (Sigma, 276685). Briefly, gelatin was dissolved at 10% (w/v) in PBS at 50 °C under constant stirring. MAA was added dropwise to the solution. The reaction proceeded for 3 h at 50 °C, after which the mixture was diluted with warm PBS, dialyzed against deionized water using 12–14 kDa MWCO tubing for 1 week at 40 °C, and lyophilized for 1 week to obtain a white solid. The degree of methacrylation was tuned by varying the MAA concentration: for low-degradability GelMA, 10 mL of MAA was added to the reaction mixture with 10 g gelatin in 100 mL solution, whereas for high-degradability GelMA, 1 mL of MAA was used under identical reaction conditions. FITC-conjugated GelMA was prepared following the same synthesis protocol described above. At the end of the 3-hour methacrylation reaction, fluorescein isothiocyanate (FITC (Sigma, 46950); 1 mg per 100 mg GelMA) was added, and the mixture was dialyzed and lyophilized. FITC-GelMA was blended with unlabeled GelMA at a 1:100 (w/w) ratio to provide fluorescence.

### Step Emulsion Microfluidic Device Preparation

Step-emulsion devices (based on modifications to published designs^70,71^) were fabricated using two-layer photolithography. A 4-inch silicon wafer (University Wafers) was dehydrated at 200 °C for 10 min, cleaned in a plasma asher at 30 W for 30 s and spin-coated with SU-8 2025 to achieve a ∼25 µm layer. After soft bake, the wafer was exposed through a transparency mask, post-baked, and developed to define the shallow step region. A second layer of SU-8 2150 was then spin-coated to achieve ∼100 µm height, followed by alignment to the first layer, UV exposure through a second mask, post-bake, and development. The final master thus contained a step transition from 25 µm to 100 µm channel height at the nozzle. To render the surface non-adhesive, the master was treated by vapor deposition of trichloro(1H,1H,2H,2H-perfluorooctyl)silane (Millipore Sigma 448931) in a desiccator for 30 min, followed by baking at 100 °C for 30 min.

PDMS (Dow, Sylgard 184) (9:1 base:curing agent) was poured over the master, degassed, and cured at 60 °C for 2 h. Devices were peeled from the master, access ports were punched, and slabs were bonded to glass slides by oxygen plasma treatment, followed by a 30 min bake at 100 °C. Channels were rendered hydrophobic by flushing with RainX, then dried at 100 °C. Prior to use, channels were primed with the continuous phase.

### Microsphere Production

The continuous oil phase consisted of light mineral oil (Fisher, O121-1) with 2.5% Span-80 (Sigma S6760) surfactant. The dispersed aqueous phase consisted of 10% (w/v) GelMA prepolymer containing 3 mg/mL lithium phenyl-2,4,6-trimethylbenzoylphosphinate (LAP) (Sigma 900889). Fluids were driven into the device using syringe pumps, and at the step geometry, uniform aqueous droplets were generated and collected into reservoirs of the continuous phase. Droplets were exposed to 365 nm UV light to initiate photopolymerization, yielding crosslinked GelMA microspheres. After curing, microspheres were first washed with PBS containing 1% Tween-20, followed by repeated centrifugation and exchange into PBS or cell culture media. The microspheres were packed by centrifugation at 5000g for 5 minutes followed by aspiration of supernatant. Same procedure was followed for production of both high-degradability and low degradability microspheres.

### Rheology

The viscoelastic and flow properties of packed GelMA microspheres were characterized using a rotational rheometer (TA Instruments Discovery HR-2) equipped with parallel plate geometry. Microspheres were pelleted by centrifugation, transferred to the lower plate, and the gap was adjusted to achieve a uniform packed layer. A solvent trap was used to minimize evaporation. Strain sweeps were performed at 10 rad/s over a strain range of 0.1–100%, frequency sweeps were conducted from 0.1–400 rad/s and flow sweeps were carried out over shear rates of 0.1–100 s⁻¹ to evaluate shear-thinning behavior.

### Hepatocyte Aggregation and in vitro Culture in Packed Microspheres

Neonatal human dermal fibroblasts (Lonza, CC2509) were maintained in FGM-2 media (Lonza CC-3132) and used between passages 4-8. RFP-NHDFs were generated by transduction with RFP Lentivirus. In short, plenti-CMV-MCS-RFP-SV-puro (Addgene, 109377) was co-transfected into HEK-293T cells together with psPAX2 and PMD2.G using Lipofectamine 3000 (Thermo Fisher Scientific, L3000015). Supernatants with assembled viruses were collected 36 and 60 hours after transfection, concentrated (Lenti-X concentrator, Takara Bio, 631232) and stored at −80°C prior to use. NHDFs were transduced in FGM-2 media with 10 µg/mL polybrene (Millipore Sigma, TR-1003-G) overnight and washed with fresh media. Transduced cells were subsequently selected in 5 µg/mL puromycin for 7 days before expansion.

Primary human hepatocytes (Thermo Fisher Scientific HU8200, HU8339, HU8412) were aggregated either alone or together with neonatal human dermal fibroblasts (NHDFs, Lonza CC2509) at a 1:1 ratio, with each aggregate containing ∼100 PHHs (PHH-only) or ∼100 PHHs and 100 NHDFs (PHH–NHDF). Aggregates were generated in custom-fabricated PDMS AggreWell arrays containing 400 µm diameter microwells and cultured for 3 days in hepatocyte medium (DMEM with L-glutamine, 70 ng/ml glucagon, 10% Fetal Bovine Serum, 1% ITS Universal Culture Supplement, 40 ng/ml dexamethasone, 15.4 mM HEPES, 1% Pen/Strep). PHHs were pre-labeled with CellTracker Deep Red (Thermo Fisher Scientific, C34565), and RFP-expressing NHDFs were used to enable longitudinal imaging of cultures.

Following aggregation, collected spheroids were resuspended with pre-packed GelMA microspheres (∼35:65, aggregate:microsphere ratio) to form a composite slurry. For in vitro cultures, the slurry was pipetted into PDMS gaskets (4 mm inner diameter) placed within well plates, allowing confinement of the slurry into defined circular wells with each gasket receiving ∼5 x 10^5^ PHHs with or without NHDFs. Cultures were maintained for up to 14 days with media exchanges every 48 h, and conditioned media was collected for albumin quantification using a commercial ELISA kit (Bethyl Laboratories). For assessment of hepatocyte function, cultures were incubated with 200 µg/mL indocyanine green (ICG) (Lumiprobe, 3059) for 15 minutes at 37 °C. Following incubation, the wells were washed with culture medium for an additional 15 minutes to remove excess dye, after which samples were imaged. Aggregate circularity and NHDF area fractions were calculated in QuPath (v0.5.1).

### In vivo Implantations

All animal procedures were performed in accordance with institutional guidelines and approved protocols. To compare bulk and microsphere integration, GelMA was formulated either as monolithic bulk gels or as injectable microsphere suspensions. Bulk constructs were prepared by photocrosslinking 10% (w/v) GelMA with 3 mg/mL LAP in PDMS molds under UV (365 nm) to yield cylindrical gels (∼5 mm diameter × ∼2.5 mm thickness). For the granular condition, GelMA microspheres were concentrated into a dense slurry and injected (∼50 µL volume/graft) into the fat pad of NSG mice under isoflurane anesthesia. Each animal received ∼5 X 10^5^ PHHs across all conditions. For initial optimization, injections were performed following surgical exposure of the fat pad to facilitate accurate placement before translation to ultrasound-guided injection and imaging.

For hepatocyte transplantation studies, three graft types were compared: (i) PHH-only aggregates, (ii) PHH-NHDF aggregates, and (iii) PHH-NHDF aggregates suspended in GelMA microspheres. For conditions without microspheres, aggregates were suspended in equal volumes of culture medium prior to injection. For experiments using microspheres with varying degradability, same volume of microspheres were used per graft between the conditions. All grafts were delivered into the fat pad under identical conditions. During injection, the microsphere slurry flowed smoothly under manual pressure and did not exhibit obvious clogging or flow instability. To minimize backflow along the needle tract, the needle was held in place for approximately 10 seconds following injection prior to withdrawal. The material consistently re-formed a localized deposit upon cessation of shear. For acellular hydrogel studies, mice received terminal intravenous injections of Texas Red–labeled 70 kDa dextran to visualize perfused vasculature. Explants were further processed for H&E and immunostaining, and vessel area fraction and nuclear infiltration were quantified using QuPath. For cell-containing grafts, functional assessment was performed by saphenous vein blood draws followed by human albumin quantification using a commercial ELISA kit (Bethyl Laboratories).

#### CYP3A4 Enzyme Activity

CYP3A4 activity was assessed using the Promega P450-Glo assay per the manufacturer’s instructions. In vitro constructs were pretreated with rifampicin (25 µM) for 48 hours, followed by incubation with the luciferin-IPA substrate in DMEM with penicillin/streptomycin for 1 hour. Conditioned media were then developed for 20 minutes, and luminescence was measured using a plate reader.

#### Urea Assay

Urea levels were measured using a commercial colorimetric kit (San Bio Labs) according to the manufacturer’s instructions. Conditioned media (10 µL) was mixed with 150 µL reagent (1/3 color, 2/3 acid), sealed, and incubated at 60–70 °C for 90 min. Plates were cooled on ice, and absorbance was measured at 540 nm with 650 nm background subtraction.

### Ultrasound-guided Injections and Imaging

Ultrasound studies were performed using a Vevo 3100 system (FUJIFILM VisualSonics) equipped with an MX550D linear array transducer (40 MHz center frequency). Mice were anesthetized with isoflurane and positioned supine on a heated stage. Real-time B-mode imaging was used to visualize needle placement and bolus appearance during injection. Ultrasound images of injected grafts, including acellular microspheres, PHH aggregates, PHH-NHDF aggregates, PHH-NHDF aggregates delivered with microspheres, and PHH-NHDF aggregates delivered with microspheres of varying degradability were acquired. Three-dimensional (3D) scans were acquired immediately following injection and at subsequent time points using motorized stepwise translation of the transducer.

For vascular imaging, nonlinear contrast-enhanced ultrasound was conducted at the study endpoint following intravenous administration of non-targeted microbubble contrast agents (Vevo MicroMarker; 200 µL continuous infusion via tail vein). Microbubble contrast signals were acquired using a 250S transducer (21 MHz center frequency). Perfusion signals within the graft region were quantified from contrast-mode images using VevoLab software.

### Tissue Clearing of Fat Pads Containing Hydrogel Implants

Explanted fat pads containing bulk or granular GelMA hydrogels were cleared using the CytoVista Tissue Clearing Kit (ThermoFisher, V11304) according to the manufacturer’s instructions. Briefly, tissues were fixed in 4% paraformaldehyde, rinsed in PBS, and incubated sequentially in the supplied clearing reagents to reduce lipid content and match refractive index. Cleared samples were imaged by confocal microscopy for visualization of FITC-labeled GelMA hydrogels and Texas Red dextran-perfused vasculature.

### In vitro Enzymatic Degradation Assay

Low- and high-degradability GelMA hydrogels were cast into PDMS gaskets (5 mm inner diameter; ∼50 µL volume) containing 0.25 mg/mL 2 MDa rhodamine-labeled dextran (Nanocs, DX2000-RB-1) to allow visualization of gel integrity. Gels were incubated at 37 °C in collagenase Type 4 (Worthington Biochemical, LS004189) at 0.05 mg/mL prepared in buffer containing 2.5% Triton X-100, 1 µM ZnCl₂, and 5 mM CaCl₂ in 50 mM Tris-HCl, pH 7.5. Control gels were incubated in same buffer lacking collagenase. Fluorescent images were acquired at different time points, and rhodamine signal intensity was quantified using QuPath to assess degradation kinetics.

### Immunostaining

Aggregates cultured within GelMA microspheres were fixed in 4% paraformaldehyde overnight at 4 °C, washed in PBS, and permeabilized with 0.1% Triton X-100 in PBS overnight at 4 °C. Nonspecific binding was blocked overnight at 4 °C in blocking buffer containing 5% normal donkey serum and 3% bovine serum albumin in PBS. Samples were then incubated overnight at 4°C with primary antibody against Arg-1 (Millipore Sigma, HPA003595, 1:400) and phalloidin (Abcam, ab176756, 1:50) to label F-actin. After extensive PBS washes, aggregates were incubated overnight at 4 °C with secondary antibodies (1:500), counterstained with Hoechst 33342 overnight at 4 °C, and mounted in antifade medium. Fluorescence imaging was performed using a Zeiss LSM 880 confocal microscope.

Explanted grafts were fixed in 4% paraformaldehyde for 24 h at room temperature, processed, and embedded in paraffin. Sections (5 µm) were deparaffinized in xylene, rehydrated through graded ethanol, and subjected to heat-induced antigen retrieval in citrate buffer (pH 6.0). Following antigen retrieval, autofluorescence was quenched by incubation in 3% hydrogen peroxide in methanol for 15 min, followed by light exposure for 2 h in the TiYO Autofluorescence Quenching System. Nonspecific binding was blocked with 5% normal donkey serum for 1 h at room temperature. Sections were incubated overnight at 4 °C with primary antibodies against Arg-1 (Millipore Sigma, HPA003595, 1:400), BSEP (Santa Cruz Biotechnology, sc-74500, 1:100), CD31 (R&D Systems, AF3628, 1:40), F4/80 (Thermo Fisher Scientific, 14-4801-82), CD86 (Thermo Fisher Scientific, PA5-88284,1:100), CD206 (R&D Systems, AF2535, 1:100), CYP3A4(Santa Cruz Biotechnology, sc-365415,1:100), and FITC (Jackson Immunoresearch, 200-002-037, 1:50) (for detection of FITC-labeled GelMA microspheres). After washing, slides were incubated with secondary antibodies (1:500) for 1 h at room temperature, counterstained with Hoechst 33342, and mounted. Fluorescence imaging was performed using a Zeiss LSM 880 confocal microscope, TissueFAXS whole-slide scanner (TissueGnostics), Pannoramic 250 Flash III whole slide scanner (3DHistech), or Keyence BZ-X800 microscope.

#### TUNEL Staining

Apoptotic cells were detected using the OneStep TUNEL Kit (Novus Biologicals) according to the manufacturer’s instructions. Tissue sections and incubated with TUNEL reaction mix. Negative controls omitted Terminal Deoxynucleotidyl Transferase (TdT), and positive controls were generated by DNase-I treatment of serial sections. TUNEL-positive cells were analyzed and quantified using QuPath.

### Statistics

All statistical analyses were performed using Prism software (GraphPad Software Inc.). Data are presented as the mean ± SD. A comparison between two groups was conducted using the unpaired Student’s t-test. Comparisons between multiple groups were performed using a one-way ANOVA test followed by Tukey’s post hoc test. Details for statistical test performed are mentioned in figure legends where applicable. For each test, p < 0.05 was considered statistically significant. For statistical significance, (*) indicates p < 0.05, (**) indicates p < 0.01, and (***) indicates p < 0.001.

## Acknowledgment

The authors would like to thank Sue Kangiser, Lian-Ee Ch’ng, and Brooks Firth Bard for providing administrative support. We also acknowledge Claire M. Zheng for her contributions, Kathleen S. Cormier for her insightful discussions, and Tuan Luo, Weija Zhang, and Charlene Condon for their assistance with histology. We would also like to acknowledge Dr. Magalie Boucher, DVM, MS, DACVP for the histopathological evaluation of the samples. The schematics were generated using BioRender. The work was supported in part by the Koch Institute Support (core) Grant P30-CA14051 from the National Cancer Institute, and we thank the Koch Institute’s Robert A. Swanson (1969) Biotechnology Center for technical support, specifically the Preclinical Imaging and Testing Core, The Hope Babette Tang (1983) Histology Facility, and the Peterson (1957) Nanotechnology Materials Core Facility. This work was supported by NIH (EB033821) and the Wellcome Leap HOPE Program. A.D.W. acknowledges support from National Science Foundation Graduate Research Fellowship. S.N.B. is a Howard Hughes Medical Institute Investigator.

## References

1. Black, C.K., Termanini, K.M., Aguirre, O., Hawksworth, J.S., and Sosin, M. (2018). Solid organ transplantation in the 21st century. Ann. Transl. Med 6, 409–409. 10.21037/atm.2018.09.68.

2. Terrault, N.A., Francoz, C., Berenguer, M., Charlton, M., and Heimbach, J. (2023). Liver Transplantation 2023: Status Report, Current and Future Challenges. Clinical Gastroenterology and Hepatology 21, 2150–2166. 10.1016/j.cgh.2023.04.005.

3. Bhatia, S.N., Underhill, G.H., Zaret, K.S., and Fox, I.J. (2014). Cell and tissue engineering for liver disease. Science Translational Medicine 6, 245sr2–245sr2. 10.1126/scitranslmed.3005975.

4. Bram, Y., Nguyen, D.-H.T., Gupta, V., Park, J., Richardson, C., Chandar, V., and Schwartz, R.E. (2021). Cell and Tissue Therapy for the Treatment of Chronic Liver Disease. Annu. Rev. Biomed. Eng. 23, 517–546. 10.1146/annurev-bioeng-112619-044026.

5. Nulty, J., Anand, H., and Dhawan, A. (2024). Human Hepatocyte Transplantation: Three Decades of Clinical Experience and Future Perspective. Stem Cells Translational Medicine 13, 204–218. 10.1093/stcltm/szad084.

6. Zhang, Y., Li, L., Dong, L., Cheng, Y., Huang, X., Xue, B., Jiang, C., Cao, Y., and Yang, J. (2024). Hydrogel-Based Strategies for Liver Tissue Engineering. Chem Bio Eng. 1, 887–915. 10.1021/cbe.4c00079.

7. Wang, Q., Feng, Y., Wang, A., Hu, Y., Cao, Y., Zheng, J., Le, Y., and Liu, J. (2024). Innovations in 3D bioprinting and biomaterials for liver tissue engineering: Paving the way for tissue-engineered liver. iLIVER 3, 100080. 10.1016/j.iliver.2024.100080.

8. Ma, X., Qu, X., Zhu, W., Li, Y.-S., Yuan, S., Zhang, H., Liu, J., Wang, P., Lai, C.S.E., Zanella, F., et al. (2016). Deterministically patterned biomimetic human iPSC-derived hepatic model via rapid 3D bioprinting. Proc. Natl. Acad. Sci. U.S.A. 113, 2206–2211. 10.1073/pnas.1524510113.

9. Lee, J.W., Choi, Y.-J., Yong, W.-J., Pati, F., Shim, J.-H., Kang, K.S., Kang, I.-H., Park, J., and Cho, D.-W. (2016). Development of a 3D cell printed construct considering angiogenesis for liver tissue engineering. Biofabrication 8, 015007. 10.1088/1758-5090/8/1/015007.

10. Daly, A.C. (2023). Granular Hydrogels in Biofabrication: Recent Advances and Future Perspectives. Adv Healthcare Materials, 2301388. 10.1002/adhm.202301388.

11. Riley, L., Schirmer, L., and Segura, T. (2019). Granular hydrogels: emergent properties of jammed hydrogel microparticles and their applications in tissue repair and regeneration. Current Opinion in Biotechnology 60, 1–8. 10.1016/j.copbio.2018.11.001.

12. Qazi, T.H., and Burdick, J.A. (2021). Granular hydrogels for endogenous tissue repair. Biomaterials and Biosystems 1, 100008. 10.1016/j.bbiosy.2021.100008.

13. Griffin, D.R., Weaver, W.M., Scumpia, P.O., Di Carlo, D., and Segura, T. (2015). Accelerated wound healing by injectable microporous gel scaffolds assembled from annealed building blocks. Nature Mater 14, 737–744. 10.1038/nmat4294.

14. Griffin, D.R., Archang, M.M., Kuan, C.-H., Weaver, W.M., Weinstein, J.S., Feng, A.C., Ruccia, A., Sideris, E., Ragkousis, V., Koh, J., et al. (2021). Activating an adaptive immune response from a hydrogel scaffold imparts regenerative wound healing. Nat. Mater. 20, 560–569. 10.1038/s41563-020-00844-w.

15. Nih, L.R., Sideris, E., Carmichael, S.T., and Segura, T. (2017). Injection of Microporous Annealing Particle (MAP) Hydrogels in the Stroke Cavity Reduces Gliosis and Inflammation and Promotes NPC Migration to the Lesion. Advanced Materials 29, 1606471. 10.1002/adma.201606471.

16. Rodriguez-Rivera, G.J., Sharma, S., Maduka, C.V., Boyd, S., Perry, A.R., Di Caprio, N., Riley, L., Miksch, C.E., Lee, D., Segura, T., et al. (2025). Microgel Aspect Ratio Influences Injectable Granular Hydrogel Scaffold Pore Structure and Cellular Invasion for Tissue Repair. Advanced Science, e11513. 10.1002/advs.202511513.

17. Hoque, J., Zeng, Y., Newman, H., Gonzales, G., Lee, C., and Varghese, S. (2022). Microgel-Assisted Delivery of Adenosine to Accelerate Fracture Healing. ACS Biomater. Sci. Eng. 8, 4863–4872. 10.1021/acsbiomaterials.2c00977.

18. Stevens, K.R., Scull, M.A., Ramanan, V., Fortin, C.L., Chaturvedi, R.R., Knouse, K.A., Xiao, J.W., Fung, C., Mirabella, T., Chen, A.X., et al. (2017). In situ expansion of engineered human liver tissue in a mouse model of chronic liver disease. Sci Transl Med 9, eaah5505. 10.1126/scitranslmed.aah5505.

19. Chen, A.A., Thomas, D.K., Ong, L.L., Schwartz, R.E., Golub, T.R., and Bhatia, S.N. (2011). Humanized mice with ectopic artificial liver tissues. Proceedings of the National Academy of Sciences 108, 11842–11847. 10.1073/pnas.1101791108.

20. Chen, A.X., Chhabra, A., Song, H.G., Fleming, H.E., Chen, C.S., and Bhatia, S.N. (2020). Controlled Apoptosis of Stromal Cells to Engineer Human Microlivers. Adv Funct Materials 30, 1910442. 10.1002/adfm.201910442.

21. March, S., Ramanan, V., Trehan, K., Ng, S., Galstian, A., Gural, N., Scull, M.A., Shlomai, A., Mota, M.M., Fleming, H.E., et al. (2015). Micropatterned coculture of primary human hepatocytes and supportive cells for the study of hepatotropic pathogens. Nat Protoc 10, 2027–2053. 10.1038/nprot.2015.128.

22. Stevens, K.R., Ungrin, M.D., Schwartz, R.E., Ng, S., Carvalho, B., Christine, K.S., Chaturvedi, R.R., Li, C.Y., Zandstra, P.W., Chen, C.S., et al. (2013). InVERT molding for scalable control of tissue microarchitecture. Nat Commun 4, 1847. 10.1038/ncomms2853.

23. Gural, N., Mancio-Silva, L., Miller, A.B., Galstian, A., Butty, V.L., Levine, S.S., Patrapuvich, R., Desai, S.P., Mikolajczak, S.A., Kappe, S.H.I., et al. (2018). In Vitro Culture, Drug Sensitivity, and Transcriptome of Plasmodium Vivax Hypnozoites. Cell Host & Microbe 23, 395–406.e4. 10.1016/j.chom.2018.01.002.

24. Kukla, D.A., Crampton, A.L., Wood, D.K., and Khetani, S.R. (2020). Microscale Collagen and Fibroblast Interactions Enhance Primary Human Hepatocyte Functions in Three-Dimensional Models. Gene Expr 20, 1–18. 10.3727/105221620X15868728381608.

25. Brown, G.E., Bodke, V.V., Ware, B.R., and Khetani, S.R. (2025). Liver portal fibroblasts induce the functions of primary human hepatocytes in vitro. Commun Biol 8, 721. 10.1038/s42003-025-08135-3.

26. Iansante, V., Mitry, R.R., Filippi, C., Fitzpatrick, E., and Dhawan, A. (2018). Human hepatocyte transplantation for liver disease: current status and future perspectives. Pediatr Res 83, 232–240. 10.1038/pr.2017.284.

27. Soltys, K.A., Soto-Gutiérrez, A., Nagaya, M., Baskin, K.M., Deutsch, M., Ito, R., Shneider, B.L., Squires, R., Vockley, J., Guha, C., et al. (2010). Barriers to the successful treatment of liver disease by hepatocyte transplantation. Journal of Hepatology 53, 769–774. 10.1016/j.jhep.2010.05.010.

28. Higashi, H., Yagi, H., Kuroda, K., Tajima, K., Kojima, H., Nishi, K., Morisaku, T., Hirukawa, K., Fukuda, K., Matsubara, K., et al. (2022). Transplantation of bioengineered liver capable of extended function in a preclinical liver failure model. American Journal of Transplantation 22, 731–744. 10.1111/ajt.16928.

29. Madhurakkat Perikamana, S.K., Seale, N., Hoque, J., Ryu, J.H., Kumar, V., Shih, Y.V., and Varghese, S. (2022). Molecularly Tailored Interface for Long-Term Xenogeneic Cell Transplantation. Adv Funct Materials 32, 2108221. 10.1002/adfm.202108221.

30. Le Guilcher, C., Merlen, G., Dellaquila, A., Labour, M.-N., Aid, R., Tordjmann, T., Letourneur, D., and Simon-Yarza, T. (2023). Engineered human liver based on pullulan-dextran hydrogel promotes mice survival after liver failure. Materials Today Bio 19, 100554. 10.1016/j.mtbio.2023.100554.

31. Lieberthal, T.J., Sahakyants, T., Szabo-Wexler, N.R., Hancock, M.J., Spann, A.P., Oliver, M.S., Grindy, S.C., Neville, C.M., and Vacanti, J.P. (2024). Implantable 3D printed hydrogels with intrinsic channels for liver tissue engineering. Proc. Natl. Acad. Sci. U.S.A. 121, e2403322121. 10.1073/pnas.2403322121.

32. Deng, B., Ma, Y., Huang, J., He, R., Luo, M., Mao, L., Zhang, E., Zhao, Y., Wang, X., Wang, Q., et al. (2024). Revitalizing liver function in mice with liver failure through transplantation of 3D-bioprinted liver with expanded primary hepatocytes. Sci. Adv. 10, eado1550. 10.1126/sciadv.ado1550.

33. Elisseeff, J., Anseth, K., Sims, D., McIntosh, W., Randolph, M., and Langer, R. (1999). Transdermal photopolymerization for minimally invasive implantation. Proc. Natl. Acad. Sci. U.S.A. 96, 3104–3107. 10.1073/pnas.96.6.3104.

34. Pleguezuelos-Beltrán, P., Nieto-García, D., Chocarro-Wrona, C., De Vicente, J., Gálvez-Martín, P., Entrena, J.M., López-Ruiz, E., and Marchal, J.A. (2025). A Novel Sprayable Fibrinogen/Glycosaminoglycans/Collagen-Based Bioink for Skin Wound Healing Applied by a Handheld Dual-Head Airbrush. Adv Healthcare Materials, e00702. 10.1002/adhm.202500702.

35. Zhao, W., Hu, C., and Xu, T. (2023). In vivo bioprinting: Broadening the therapeutic horizon for tissue injuries. Bioactive Materials 25, 201–222. 10.1016/j.bioactmat.2023.01.018.

36. Davoodi, E., Li, J., Ma, X., Najafabadi, A.H., Yoo, J., Lu, G., Sani, E.S., Lee, S., Montazerian, H., Kim, G., et al. (2025). Imaging-guided deep tissue in vivo sound printing. Science 388, 616–623. 10.1126/science.adt0293.

37. Correa, S., Grosskopf, A.K., Lopez Hernandez, H., Chan, D., Yu, A.C., Stapleton, L.M., and Appel, E.A. (2021). Translational Applications of Hydrogels. Chem. Rev. 121, 11385–11457. 10.1021/acs.chemrev.0c01177.

38. Lim, K.S., Klotz, B.J., Lindberg, G.C.J., Melchels, F.P.W., Hooper, G.J., Malda, J., Gawlitta, D., and Woodfield, T.B.F. (2019). Visible Light Cross-Linking of Gelatin Hydrogels Offers an Enhanced Cell Microenvironment with Improved Light Penetration Depth. Macromol Biosci 19, e1900098. 10.1002/mabi.201900098.

39. Zennifer, A., Manivannan, S., Sethuraman, S., Kumbar, S.G., and Sundaramurthi, D. (2022). 3D bioprinting and photocrosslinking: emerging strategies & future perspectives. Biomater Adv 134, 112576. 10.1016/j.msec.2021.112576.

40. Zhang, H., Tang, B., Zhang, B., Huang, K., Li, S., Zhang, Y., Zhang, H., Bai, L., Wu, Y., Cheng, Y., et al. (2024). X-ray-activated polymerization expanding the frontiers of deep-tissue hydrogel formation. Nat Commun 15, 3247. 10.1038/s41467-024-47559-z.

41. Jaberi, A., Ghelich, P., Samandari, M., Kheirabadi, S., Ataie, Z., Kedzierski, A., Hassani Najafabadi, A., Tamayol, A., and Sheikhi, A. (2025). Gelatin methacryloyl granular hydrogel scaffolds for skin wound healing. Biomater. Sci., 10.1039.D4BM01062K. 10.1039/D4BM01062K.

42. Hsu, C.-C., George, J.H., Waller, S., Besnard, C., Nagel, D.A., Hill, E.J., Coleman, M.D., Korsunsky, A.M., Cui, Z., and Ye, H. (2022). Increased connectivity of hiPSC-derived neural networks in multiphase granular hydrogel scaffolds. Bioactive Materials 9, 358–372. 10.1016/j.bioactmat.2021.07.008.

43. Mohindra, P., Zhong, J.X., Fang, Q., Cuylear, D.L., Huynh, C., Qiu, H., Gao, D., Kharbikar, B.N., Huang, X., Springer, M.L., et al. (2023). Local decorin delivery via hyaluronic acid microrods improves cardiac performance, ventricular remodeling after myocardial infarction. npj Regen Med 8, 60. 10.1038/s41536-023-00336-w.

44. Caprio, N.D., Davidson, M.D., Daly, A.C., and Burdick, J.A. (2024). Injectable MSC Spheroid and Microgel Granular Composites for Engineering Tissue. Advanced Materials 36, 2312226. 10.1002/adma.202312226.

45. Zhu, Y., Sun, Y., Rui, B., Lin, J., Shen, J., Xiao, H., Liu, X., Chai, Y., Xu, J., and Yang, Y. (2022). A Photoannealed Granular Hydrogel Facilitating Hyaline Cartilage Regeneration via Improving Chondrogenic Phenotype. ACS Appl. Mater. Interfaces 14, 40674–40687. 10.1021/acsami.2c11956.

46. Song, W., Ma, Z., Wang, X., Wang, Y., Wu, D., Wang, C., He, D., Kong, L., Yu, W., Li, J.J., et al. (2023). Macroporous Granular Hydrogels Functionalized with Aligned Architecture and Small Extracellular Vesicles Stimulate Osteoporotic Tendon-To-Bone Healing. Advanced Science 10, 2304090. 10.1002/advs.202304090.

47. Zoratto, N., Di Lisa, D., De Rutte, J., Sakib, M.N., Alves E Silva, A.R., Tamayol, A., Di Carlo, D., Khademhosseini, A., and Sheikhi, A. (2020). In situ forming microporous gelatin methacryloyl hydrogel scaffolds from thermostable microgels for tissue engineering. Bioengineering & Transla Med 5, e10180. 10.1002/btm2.10180.

48. Sheikhi, A., De Rutte, J., Haghniaz, R., Akouissi, O., Sohrabi, A., Di Carlo, D., and Khademhosseini, A. (2019). Microfluidic-enabled bottom-up hydrogels from annealable naturally-derived protein microbeads. Biomaterials 192, 560–568. 10.1016/j.biomaterials.2018.10.040.

49. Caldwell, A.S., Aguado, B.A., and Anseth, K.S. (2020). Designing Microgels for Cell Culture and Controlled Assembly of Tissue Microenvironments. Adv Funct Materials 30, 1907670. 10.1002/adfm.201907670.

50. Caldwell, A.S., Campbell, G.T., Shekiro, K.M.T., and Anseth, K.S. (2017). Clickable Microgel Scaffolds as Platforms for 3D Cell Encapsulation. Adv Healthcare Materials 6, 1700254. 10.1002/adhm.201700254.

51. Fox, I. (2004). Hepatocyte transplantation. Journal of Hepatology 40, 878–886. 10.1016/j.jhep.2004.04.009.

52. Chen, C., Soto-Gutierrez, A., Baptista, P.M., and Spee, B. (2018). Biotechnology Challenges to In Vitro Maturation of Hepatic Stem Cells. Gastroenterology 154, 1258–1272. 10.1053/j.gastro.2018.01.066.

53. Zhang, K., Zhang, L., Liu, W., Ma, X., Cen, J., Sun, Z., Wang, C., Feng, S., Zhang, Z., Yue, L., et al. (2018). In Vitro Expansion of Primary Human Hepatocytes with Efficient Liver Repopulation Capacity. Cell Stem Cell 23, 806–819.e4. 10.1016/j.stem.2018.10.018.

54. Zhang, K., Yuan, X., Lu, S., Shu, Y., Wang, C., Cen, J., Wu, B., and Hui, L. (2025). Expansion of human hepatocytes and their application in three-dimensional culture and genetic manipulation. Nat Protoc. 10.1038/s41596-025-01211-2.

55. Hendriks, D., Artegiani, B., Hu, H., Chuva De Sousa Lopes, S., and Clevers, H. (2021). Establishment of human fetal hepatocyte organoids and CRISPR–Cas9-based gene knockin and knockout in organoid cultures from human liver. Nat Protoc 16, 182–217. 10.1038/s41596-020-00411-2.

56. Mallanna, S.K., Karanth, S.S., Marturano, J.E., Kudva, A.K., Lehmann, M., Morse, J.K., Jamiel, M., Norman, T., Wilson, C., Munarin, F., et al. (2024). Expandable, Functional Hepatocytes Derived from Primary Cells Enable Liver Therapeutics. Preprint, 10.1101/2024.12.28.630269 10.1101/2024.12.28.630269.

57. Michailidis, E., Vercauteren, K., Mancio-Silva, L., Andrus, L., Jahan, C., Ricardo-Lax, I., Zou, C., Kabbani, M., Park, P., Quirk, C., et al. (2020). Expansion, in vivo–ex vivo cycling, and genetic manipulation of primary human hepatocytes. Proc. Natl. Acad. Sci. U.S.A. 117, 1678–1688. 10.1073/pnas.1919035117.

58. Feng, S., Chen, K., and Wang, S. (2025). Practical Guide to the Design of Granular Hydrogels for Customizing Complex Cellular Microenvironments. Adv Healthcare Materials, e01947. 10.1002/adhm.202501947.

59. Coronel, M.M., Martin, K.E., Hunckler, M.D., Kalelkar, P., Shah, R.M., and García, A.J. (2022). Hydrolytically Degradable Microgels with Tunable Mechanical Properties Modulate the Host Immune Response. Small 18, 2106896. 10.1002/smll.202106896.

60. Xin, S., Gregory, C.A., and Alge, D.L. (2020). Interplay between degradability and integrin signaling on mesenchymal stem cell function within poly(ethylene glycol) based microporous annealed particle hydrogels. Acta Biomaterialia 101, 227–236. 10.1016/j.actbio.2019.11.009.

61. Butenko, S., Nagalla, R.R., Guerrero-Juarez, C.F., Palomba, F., David, L.-M., Nguyen, R.Q., Gay, D., Almet, A.A., Digman, M.A., Nie, Q., et al. (2024). Hydrogel crosslinking modulates macrophages, fibroblasts, and their communication, during wound healing. Nat Commun 15, 6820. 10.1038/s41467-024-50072-y.

62. Zhuang, Z., Zhang, Y., Sun, S., Li, Q., Chen, K., An, C., Wang, L., Van Den Beucken, J.J.J.P., and Wang, H. (2020). Control of Matrix Stiffness Using Methacrylate–Gelatin Hydrogels for a Macrophage-Mediated Inflammatory Response. ACS Biomater. Sci. Eng. 6, 3091–3102. 10.1021/acsbiomaterials.0c00295.

63. Liu, Y., Suarez-Arnedo, A., Shetty, S., Wu, Y., Schneider, M., Collier, J.H., and Segura, T. (2023). A Balance between Pro-Inflammatory and Pro-Reparative Macrophages is Observed in Regenerative D-MAPS. Adv Sci (Weinh) 10, e2204882. 10.1002/advs.202204882.

64. Sadtler, K., Estrellas, K., Allen, B.W., Wolf, M.T., Fan, H., Tam, A.J., Patel, C.H., Luber, B.S., Wang, H., Wagner, K.R., et al. (2016). Developing a pro-regenerative biomaterial scaffold microenvironment requires T helper 2 cells. Science 352, 366–370. 10.1126/science.aad9272.

65. Hui, E.E., and Bhatia, S.N. (2007). Micromechanical control of cell–cell interactions. Proc. Natl. Acad. Sci. U.S.A. 104, 5722–5726. 10.1073/pnas.0608660104.

66. Westerfield, A.D., Grzelak, K.A., Katsuyama, K., Kumar, V., Miller, B.M., Yun, J., Kirkpatrick, J., Mankus, D., Bisher, M.E., Lytton-Jean, A.K.R., et al. (2025). A 3D in vitro model of the human hepatobiliary junction. Preprint, 10.1101/2025.07.11.664464 10.1101/2025.07.11.664464.

67. Deo, K.A., Murali, A., Tronolone, J.J., Mandrona, C., Lee, H.P., Rajput, S., Hargett, S.E., Selahi, A., Sun, Y., Alge, D.L., et al. (2024). Granular Biphasic Colloidal Hydrogels for 3D Bioprinting. Adv Healthcare Materials, 2303810. 10.1002/adhm.202303810.

68. Seymour, A.J., Shin, S., and Heilshorn, S.C. (2021). 3D Printing of Microgel Scaffolds with Tunable Void Fraction to Promote Cell Infiltration. Adv Healthcare Materials 10, 2100644. 10.1002/adhm.202100644.

69. Basu, L., Bhagat, V., Ching, M.E.A., Di Giandomenico, A., Dostie, S., Greenberg, D., Greenberg, M., Hahm, J., Hilton, N.Z., Lamb, K., et al. (2023). Recent Developments in Islet Biology: A Review With Patient Perspectives. Canadian Journal of Diabetes 47, 207–221. 10.1016/j.jcjd.2022.11.003.

70. Ofner, A., Moore, D.G., Rühs, P.A., Schwendimann, P., Eggersdorfer, M., Amstad, E., Weitz, D.A., and Studart, A.R. (2017). High-Throughput Step Emulsification for the Production of Functional Materials Using a Glass Microfluidic Device. Macro Chemistry & Physics 218, 1600472. 10.1002/macp.201600472.

71. De Rutte, J.M., Koh, J., and Di Carlo, D. (2019). Scalable High-Throughput Production of Modular Microgels for In Situ Assembly of Microporous Tissue Scaffolds. Adv Funct Materials 29, 1900071. 10.1002/adfm.201900071.

