## Supplementary material for "Image-Guided Injectable Niche for Hepatocyte Transplantation": SI

**Figure S1**

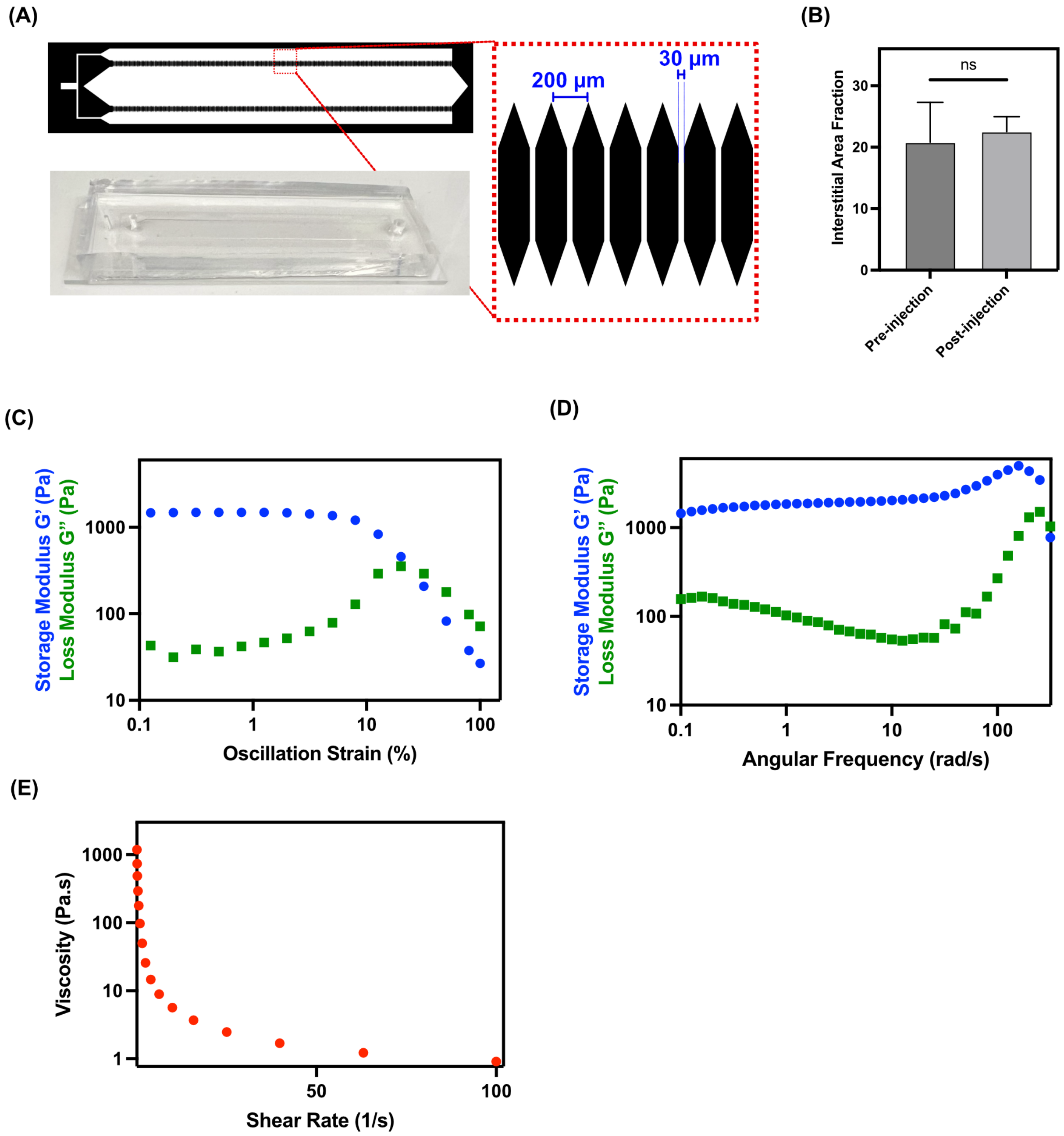
(A)
200 µm
30 µm
(B)
ns
Interstitial Area Fraction
Pre-injection
Post-injection
(C)
Storage Modulus G' (Pa)
Loss Modulus G" (Pa)
Oscillation Strain (%)
(D)
Storage Modulus G' (Pa)
Loss Modulus G" (Pa)
Angular Frequency (rad/s)
(E)
Viscosity (Pa.s)
Shear Rate (1/s)

**Figure S2**

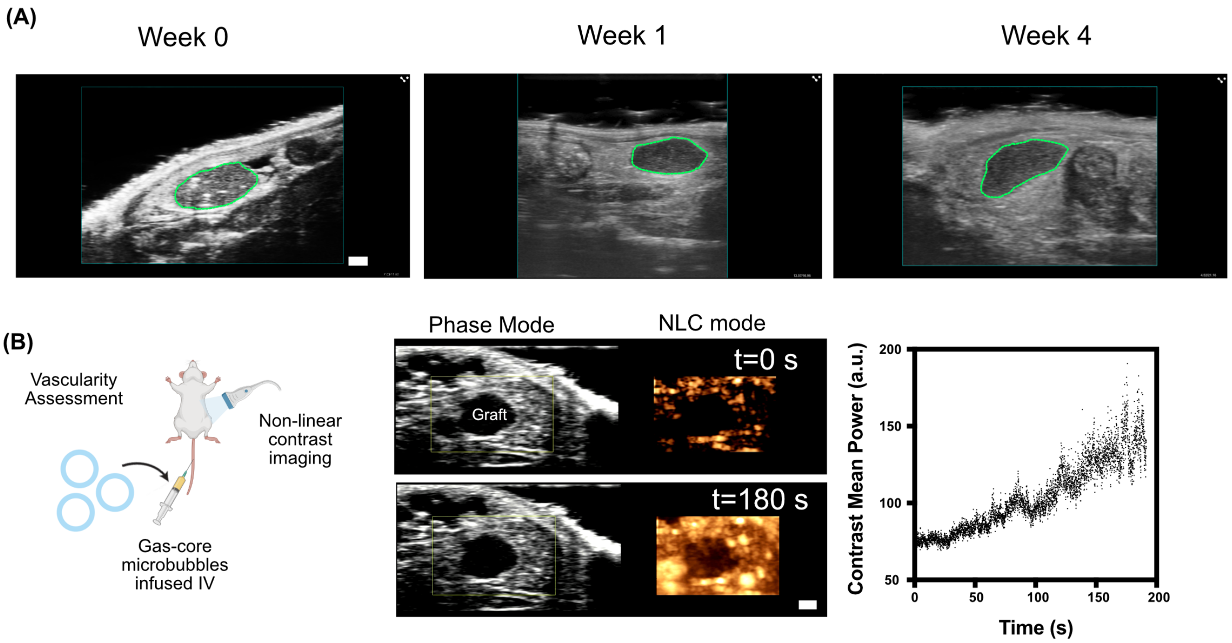

(A)
Week 0
Week 1
Week 4
(B)
Vascularity Assessment
Non-linear contrast imaging
Gas-core microbubbles infused IV
Phase Mode
NLC mode
t=0 s
Graft
t=180 s
Contrast Mean Power (a.u.)
200
150
100
50
0
50
100
150
200
Time (s)

**Figure S3**

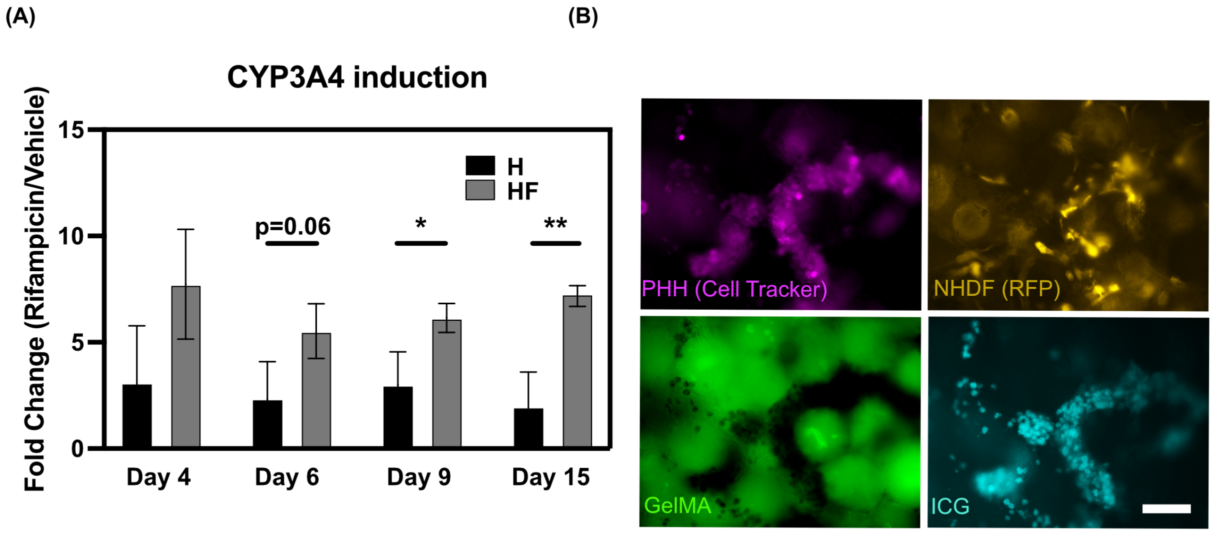

(A)
CYP3A4 induction
Fold Change (Rifampicin/Vehicle)
15
10
5
0
H
HF
p=0.06
*
**
Day 4
Day 6
Day 9
Day 15
(B)
PHH (Cell Tracker)
NHDF (RFP)
GelMA
ICG

#### Figure S4

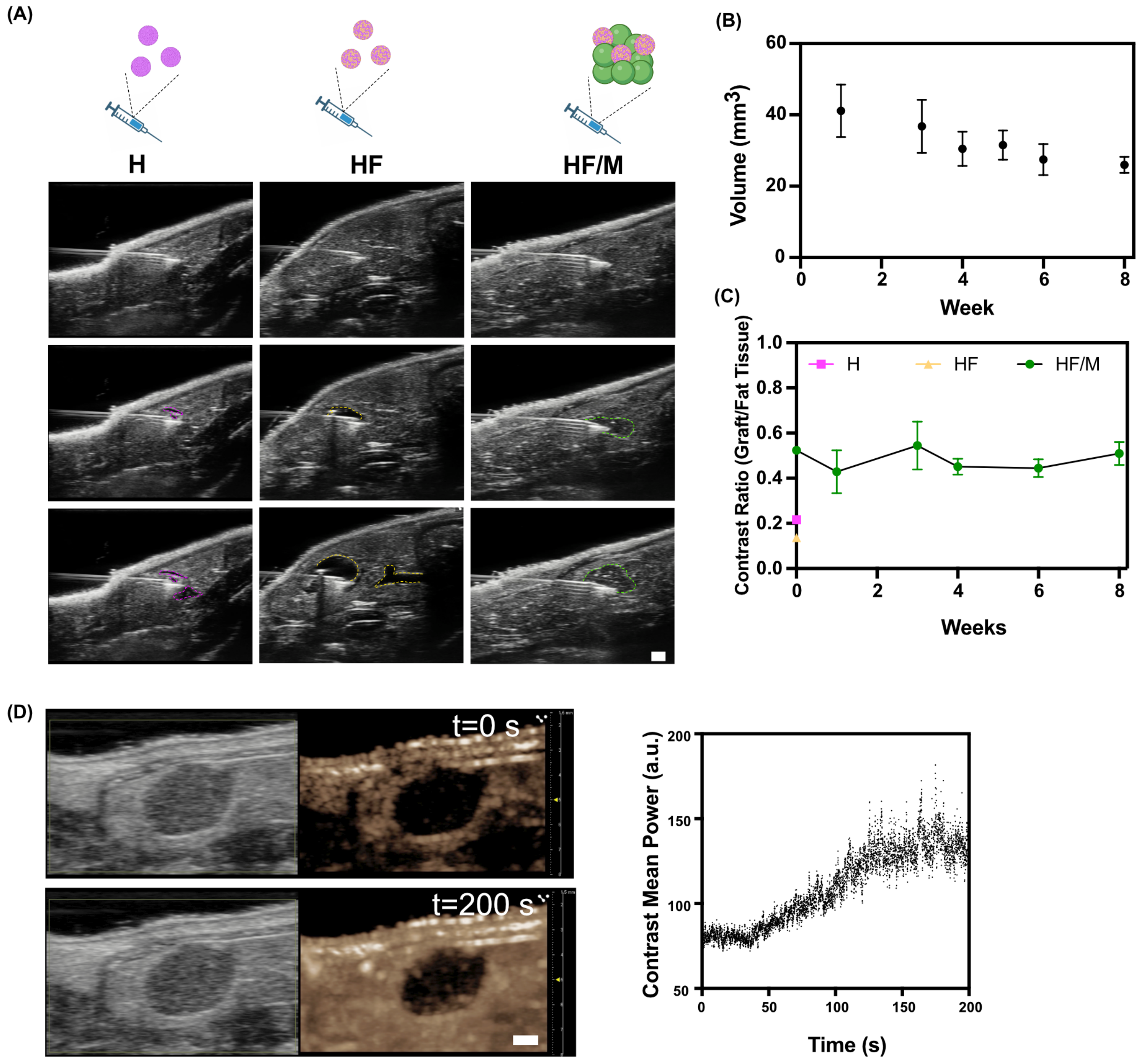

(A)
H
HF
HF/M
(B)
Volume (mm3)
Week
(C)
Contrast Ratio (Graft/Fat Tissue)
H
HF
HF/M
Weeks
(D)
t=0 s
t=200 s
Contrast Mean Power (a.u.)
Time (s)

**Figure S5**

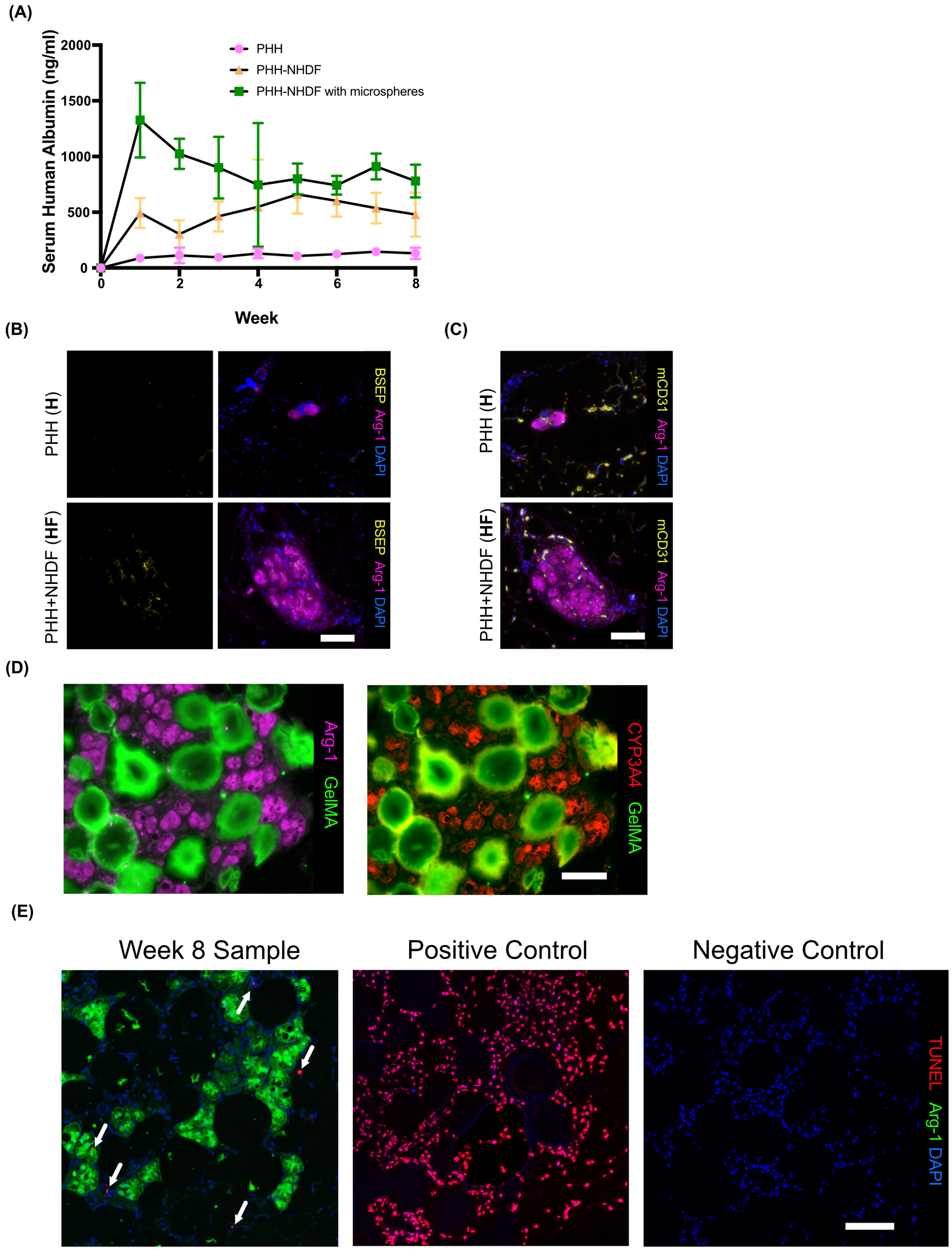
(A)
Serum Human Albumin (ng/ml)
2000
1500
1000
500
0
0
2
4
6
8
Week
PHH
PHH-NHDF
PHH-NHDF with microspheres
(B)
PHH (H)
PHH+NHDF (HF)
BSEP Arg-1 DAPI
(C)
PHH (H)
PHH+NHDF (HF)
mCD31 Arg-1 DAPI
(D)
Arg-1 GelMA
CYP3A4 GelMA
(E)
Week 8 Sample
Positive Control
Negative Control
TUNEL Arg-1 DAPI

### Figure S6

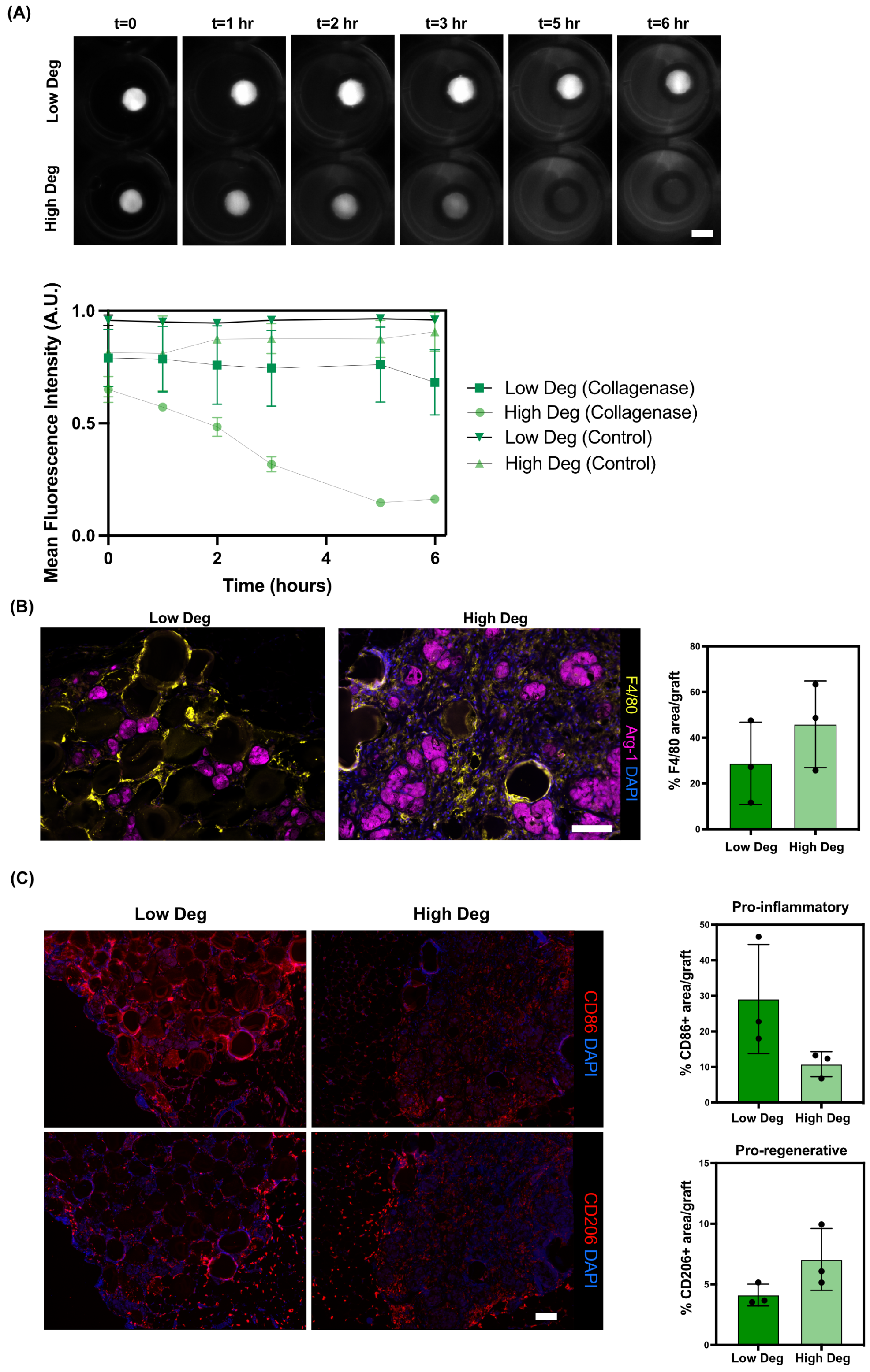
(A)
t=0
t=1 hr
t=2 hr
t=3 hr
t=5 hr
t=6 hr
Low Deg
High Deg
Mean Fluorescence Intensity (A.U.)
1.0
0.5
0.0
0
2
4
6
Time (hours)
Low Deg (Collagenase)
High Deg (Collagenase)
Low Deg (Control)
High Deg (Control)
(B)
Low Deg
High Deg
F4/80 Arg-1 DAPI
% F4/80 area/graft
80
60
40
20
0
Low Deg
High Deg
(C)
Low Deg
High Deg
CD86 DAPI
CD206 DAPI
Pro-inflammatory
% CD86+ area/graft
50
40
30
20
10
0
Low Deg
High Deg
Pro-regenerative
% CD206+ area/graft
15
10
5
0
Low Deg
High Deg

**Figure S1.** Characterization of GelMA microspheres and rheological properties. (A) Step-emulsification device design showing nozzle architecture (top), fabricated PDMS device (bottom), and magnified channel dimensions. (B) Quantification of interstitial area fraction pre- and post-injection demonstrating preservation of porosity (n=6 and n=5 samples for pre-injection and post-injection respectively). (C) Representative strain sweep showing storage (G′) and loss (G″) moduli of packed GelMA microspheres (n=3). (D) Representative frequency sweep demonstrating viscoelastic behavior (n=3). (E) Representative shear-thinning behavior as measured by viscosity versus shear rate (n=2).

**Figure S2.** Longitudinal ultrasound imaging of graft persistence and vascular connectivity. (A) Representative B-mode ultrasound images showing persistence of echogenic acellular GelMA microsphere grafts at weeks 0, 1, and 4 post-injection. (B) Schematic of vascularity assessment using intravenous microbubble injection and nonlinear contrast (NLC) imaging. Phase and NLC mode images of graft regions at baseline (t = 0 s) and maximum intensity projection after 180 s demonstrate progressive contrast enhancement. Right: quantification of mean contrast power over time confirms microbubble perfusion within the graft (Scale bars 1 mm).

**Figure S3.** In vitro functional characterization of PHHs. (A) CYP3A4 activity assessed by rifampicin induction and expressed as fold change relative to vehicle control at the indicated time points. (n=3 in vitro constructs for each condition). (B) Representative images showing PHHs labeled with CellTracker (magenta), NHDFs expressing RFP (yellow), GelMA microspheres (green), and hepatocyte-specific indocyanine green (ICG) uptake (cyan) (Scale bar 100 μm). For statistical significance, (*) indicates $p < 0.05$ and (**) indicates $p < 0.01$.

**Figure S4.** Ultrasound-guided injection and monitoring of microspheres-based liver grafts. (A) Representative B-mode images showing injection of PHH aggregates alone (top) PHH-NHDF aggregates alone (middle) versus aggregates with GelMA microspheres (bottom), with graft outlines indicated. (Scale bar 1 mm) (B) Quantification of graft volume over time (mean ± SD). (C) Contrast ratio of graft to surrounding fat pad over time, comparing microsphere-containing grafts to aggregates alone. (D) Representative nonlinear contrast-enhanced ultrasound at baseline (t = 0 s) and maximum intensity projection image at 200s into microbubble injection, showing progressive perfusion of the graft region. Right: quantification of mean contrast power over time within the graft region (Scale bar 1 mm).

**Figure S5.** Functional and histological characterization of graft. (A) Longitudinal measurement of human albumin levels in serum following transplantation of PHHs alone, PHH–NHDF co-cultures, or PHH–NHDF constructs incorporating microspheres, measured over 8 weeks. (B) Representative immunofluorescence images of explanted grafts stained for the bile salt export pump (BSEP, yellow) and Arg-1 (magenta), with nuclei counterstained with DAPI (blue) for PHHs alone (H) and PHH–NHDF (HF) conditions. (C) Immunofluorescence staining for mouse CD31 (yellow) and Arg-1 (magenta) demonstrating host-derived vascularization within PHH and PHH–NHDF grafts. (Scale bar 100 μm) (D) CYP3A4 immunostaining (red) for grafts containing PHH–

NHDF aggregates in GelMA microspheres (Scale bar 100 μm). (E) Representative images of TUNEL staining in sections of explanted graft at 8 weeks. White arrows point to TUNEL+ stain within the image. Positive controls were generated by treating a consecutive tissue section with DNase I prior to staining, which induces DNA strand breaks and confirms that the TUNEL assay can detect damaged DNA. Negative controls were generated by omitting the Terminal Deoxynucleotidyl Transferase (TdT) enzyme during the assay; the absence of signal in these sections demonstrates that staining is enzyme-dependent with low background. For direct comparison, the same region of the graft is shown across experimental and control conditions using serial sections (Scale bar 100 μm).

**Figure S6.** Degradability of GelMA microspheres influences remodeling and immune cell distribution. (A) In vitro enzymatic degradation of low- and high-degradability hydrogels assessed by fluorescence intensity over 6 h. (B) Immunofluorescence staining and quantification of explanted grafts at 4 weeks showing hepatocytes (Arg-1, magenta), macrophages (F4/80, yellow), and nuclei (blue) in low- versus high-degradability microsphere conditions (Scale bar 100 μm). (C) Representative immunofluorescence images and quantification of explanted grafts stained for CD86 (red, top) and CD206 (red, bottom). For each sample, >6 ROIs per graft were stitched and analyzed (n=3 animals per group) (Scale bar 100 μm).
